# Combined multidimensional single-cell protein and RNA profiling dissects the cellular and functional heterogeneity of thymic epithelial cells

**DOI:** 10.1101/2022.09.14.507949

**Authors:** Fabian Klein, Clara Veiga-Villauriz, Anastasiya Börsch, Stefano Maio, Sam Palmer, Saulius Zuklys, Irene Calvo-Asensio, Lucas Musette, Mary E. Deadman, Fatima Dhalla, Andrea White, Beth Lucas, Graham Anderson, Georg A. Holländer

## Abstract

The network of thymic stromal cells provides essential niches with unique molecular cues controlling T-cell development and selection. Recent single-cell RNA-sequencing studies uncovered a large transcriptional heterogeneity among thymic epithelial cells (TEC) demonstrating a previously unappreciated complexity. However, there are only very few cell markers that allow a comparable phenotypic identification of TEC. Here we deconvoluted by massively parallel flow cytometry and machine learning known and novel TEC phenotypes into novel subpopulations and related these by CITEseq to the corresponding TEC subtypes defined by the cells’ individual RNA profiles. This approach phenotypically identified perinatal cTEC, physically located these cells within the cortical stromal scaffold, displayed their dynamic change during the life course and revealed their exceptional efficiency in positively selecting immature thymocytes. Collectively, we have identified novel markers that allow for an unprecedented dissection of the thymus stromal complexity, the cells physical isolation and assignment of specific functions to individual TEC subpopulations.

## Introduction

The thymus is essential for the formation and maintenance of the adaptive immune system. as its stroma provides a unique microenvironment promoting the generation and selection of T lymphocytes tolerant to an individual’s own tissue antigens yet responsive to an unlimited range of pathogens or malignantly transformed cells. Thymic epithelial cells (TEC) constitute the major cellular element of the stromal scaffold (Anderson et al., 1993; Anderson and Takahama, 2012; James et al., 2021). Other cellular components of the stroma are different mesenchymal cell types and endothelial cells (Handel et al., 2022; James et al., 2021). TEC attract blood-borne lymphoid progenitors, commit them to a T cell fate, provide the molecular cues essential for expansion and differentiation, and shape the T cell antigen receptor (TCR) repertoire via stringent processes of positive and negative selection based on the cells’ antigen specificity (Kadouri et al., 2020; Klein et al., 2014; Zlotoff and Bhandoola, 2011).

The TEC compartment is composed of separate cortical (c) and medullary (m) lineages which have characteristically been defined by the cells’ anatomical location, a limited number of phenotypic markers and several functional characteristics (Abramson and Anderson, 2017; Derbinski et al., 2001; Laufer et al., 1996). The surface markers Ly51 and reactivity to UEA1 have typically been used to distinguish between cTEC (Ly51^+^UEA1^-^) and mTEC (Ly51^-^UEA1^+^). Markers such as CD80 and MHCII have further been used to identify subsets of mTEC such as immature (CD80^lo^MHCII^lo^; mTEC^lo^) and mature epithelia (CD80^hi^MHCII^hi^; mTEC^hi^). The latter cells are further differentiated based on the cells’ capacity to express the Autoimmune Regulator (Aire) (Gray et al., 2006; Herzig et al., 2017). Recent single-cell RNA-sequencing (scRNAseq) uncovered a remarkable TEC heterogeneity which could previously not be appreciated using the few cell surface markers available for flow cytometry (Baran-Gale et al., 2020; Bornstein et al., 2018; Dhalla et al., 2020). For example, a scRNAseq analysis of the TEC compartment of 4-week-old mice demonstrated a single cortical TEC type but 4 separate mTEC subtypes, namely immature and a mature mTEC, post-Aire mTEC and tuft-like mTEC (Bornstein et al., 2018). Investigations of TEC heterogeneity across the life trajectory (1 to 52-weeks of age) identified 9 different TEC subtypes whose relative frequencies vary with age (Baran-Gale et al., 2020). However, only few of these transcriptionally defined TEC subsets can currently be assigned to any cytometrically characterised TEC subpopulation. (For clarity, we refer to transcriptionally defined TEC clusters as subtypes and cytometrically specified TEC as subpopulations). This limitation hinders the isolation and functional characterization of specific TEC subtypes with known RNA expression profiles.

To address this limitation, we sought to screen mouse TEC for the expression of 260 cell surface markers employing massively parallel flow cytometry and the Infinity Flow computational pipeline to infer a co-expression pattern for any of the tested epitopes (Becht et al., 2021). This approach identified several novel TEC surface markers that when suitably combined identified perinatal cTEC, intertypical TEC and tuft-like mTEC which had previously only been classified either by their distinct RNA expression profiles or a combination of cell surface and intracellular markers (Lucas et al., 2020; Miller et al., 2018). The identity of these phenotypically defined TEC subpopulations was verified by scRNAseq and, in the case of perinatal cTEC, further characterised functionally, spatially, and developmentally.

## Results

### Establishment of a cell surface expression atlas across thymic stromal cell subsets

To resolve thymic stroma heterogeneity at a phenotypic level, we sought to identify new cell surface markers that reliably and accurately identify TEC subsets hitherto only defined by the cells’ individual gene expression profiles. For this purpose, we used massively parallel flow cytometry for 260 individual cell surface markers followed by an analysis employing machine learning to compute possible co-expression patterns. Thymic stomal cells were isolated as single cells from 1-, 4-, and 16-week-old mice, physically enriched and subsequently stained for 12 backbone markers that either alone or in combination reliably identified haematopoietic (CD45), different epithelial (EpCAM1, Ly51, UEA1, MHCII, CD40, CD80, CD86, Sca1, AIRE, Podoplanin), endothelial (CD31) and some mesenchymal cells (Sca1, Ly51, Podoplanin; Figure 1A). In a next step, each of these cells were stained separately with individual antibodies specific for any of the 260 exploratory markers. Infinity Flow, a computational machine learning algorithm based on the non-linear detection of the backbone markers, was subsequently used to impute at single-cell level the co-expression of individual exploratory markers (Becht et al., 2021). The resulting heterogeneity of phenotypes was then visualised and interpreted by the single-cell analysis pipeline, Seurat (Hao et al., 2021). This hierarchical clustering of the data resulted in 7 clusters for data drawn from 1-week-old mice and in 10 clusters for that of older animals, as illustrated in two dimensions by a Uniform Manifold Approximation and Projection (UMAP) (Figure S1A-C).

**Figure 1.**
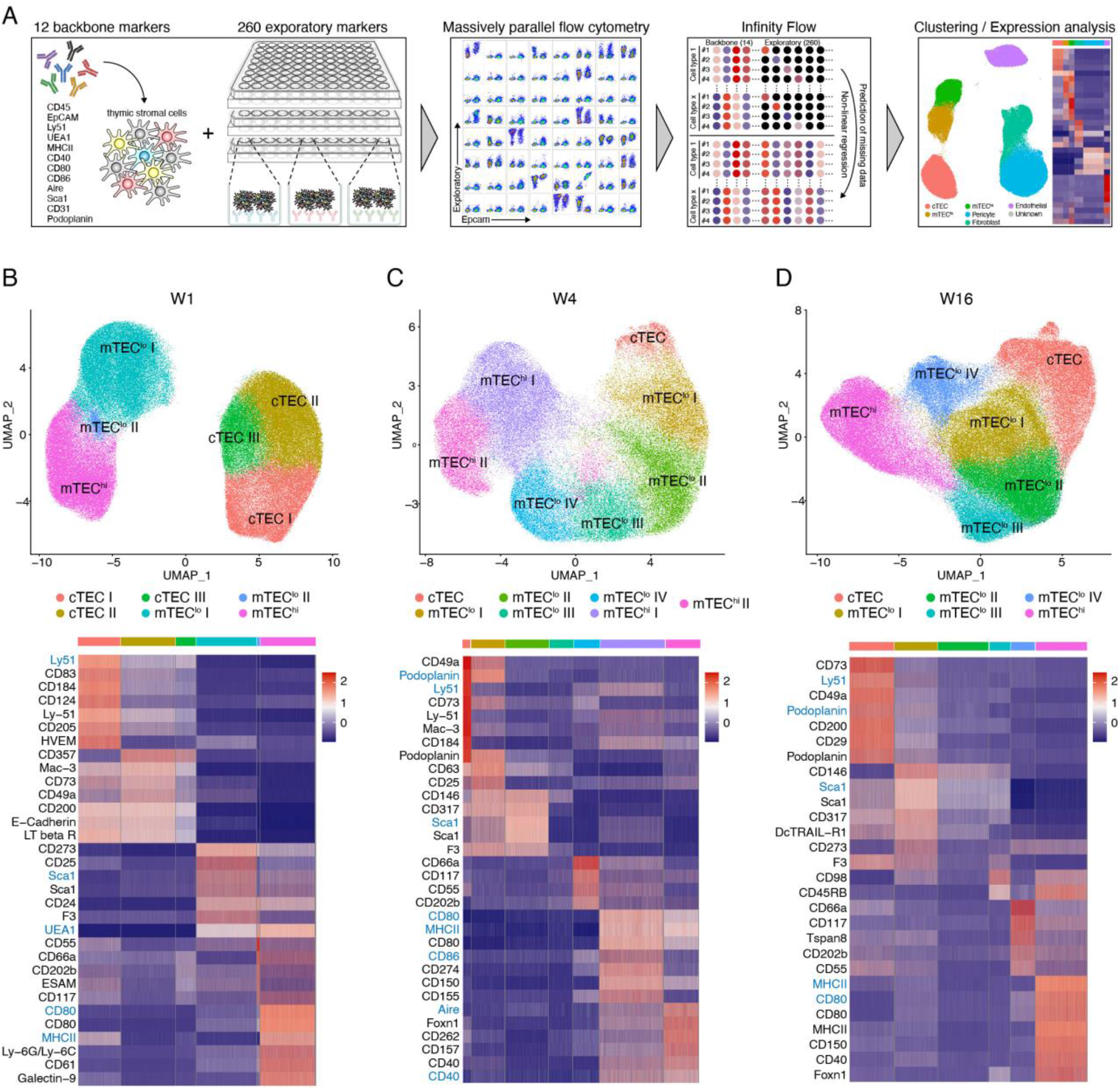
Infinity Flow analysis reveals TEC heterogeneity. **(A)** Schematic illustration of the surface marker screening pipeline. **(B-D)** Infinity Flow analysis was used to impute the expression of surface markers on TEC (CD45^-^EpCAM1^+^) derived from thymi of 1-, (C) 4-, and (D) 16-week-old mice. Hierarchical clustering analysis was performed on (B) 182123, (C) 92402, and (D) 183124 TEC, respectively, and projected in a 2-dimensional space using UMAP (top panels; 6 to 7 clusters were obtained per timepoint). Each colour represents a specific cluster as indicated. Heatmaps (bottom panels) display the expression of the top 7 markers upregulated in each cluster (log fold-change > 0.2). Backbone markers have a blue font.

At each of the three separate timepoints, the major thymic stromal cell types, epithelia, fibroblasts, pericytes, and endothelial cells, could reliably be identified based on the expression of key markers including EpCAM1 (CD326) identifying TEC, CD140a and Podoplanin marking fibroblasts, Ly51 and CD146 singling out pericytes, and CD31 staining endothelial cells. Additional markers identified subsets within these cell populations (see Figures S1A-C). Several of the antibody specificities to detect backbone epitopes were also included among the selected 260 exploratory markers (e.g. EpCAM1, CD31, Ly51, and Sca1) which allowed direct comparisons between exploratory and identical backbone markers, thus verifying the utility of the Infinity Flow algorithm. For these markers we noted identical expression profiles, therefore demonstrating the reliability of the computational approach taken (Figure S1D).

The initial expression analysis not only confirmed by flow cytometry the heterogeneity among thymic stromal cell types, but also revealed a dynamic change over time in the relative representation of individual TEC subpopulations (Figure S1A-C). A second analysis focused exclusively on EpCAM1^+^ cells and disclosed in 1-week-old but not older mice three separate cTEC subclusters as defined by the cells’ differential expression of Ly51, UEA1, MHCII, and CD80, thus illustrating a greater heterogeneity of the cTEC population early in life (Figure 1B-D and Figure S2A-C). In mice 4 weeks of age and older, mTEC with a low surface expression of MHCII (designated mTEC^lo^) segregated into 4 separate subclusters based on the differential expression of the surface markers analysed (Figure 1B-D).

### cTEC heterogeneity identified by differential cell surface marker expression

We next queried whether the expression of CD83, CD40, HVEM (CD270), and Ly51 unequivocally classified individual cTEC subpopulations, since their intensity profile differed across cTEC clusters identified in 1-week-old mice (Figure 2A). The expression of CD40 and HVEM were exclusively restricted to a subcluster designated cTEC I (see below) whereas the two other markers were detected across all cTEC subclusters, but with a stronger signal on the cTEC I (Figure 2A).

**Figure 2.**
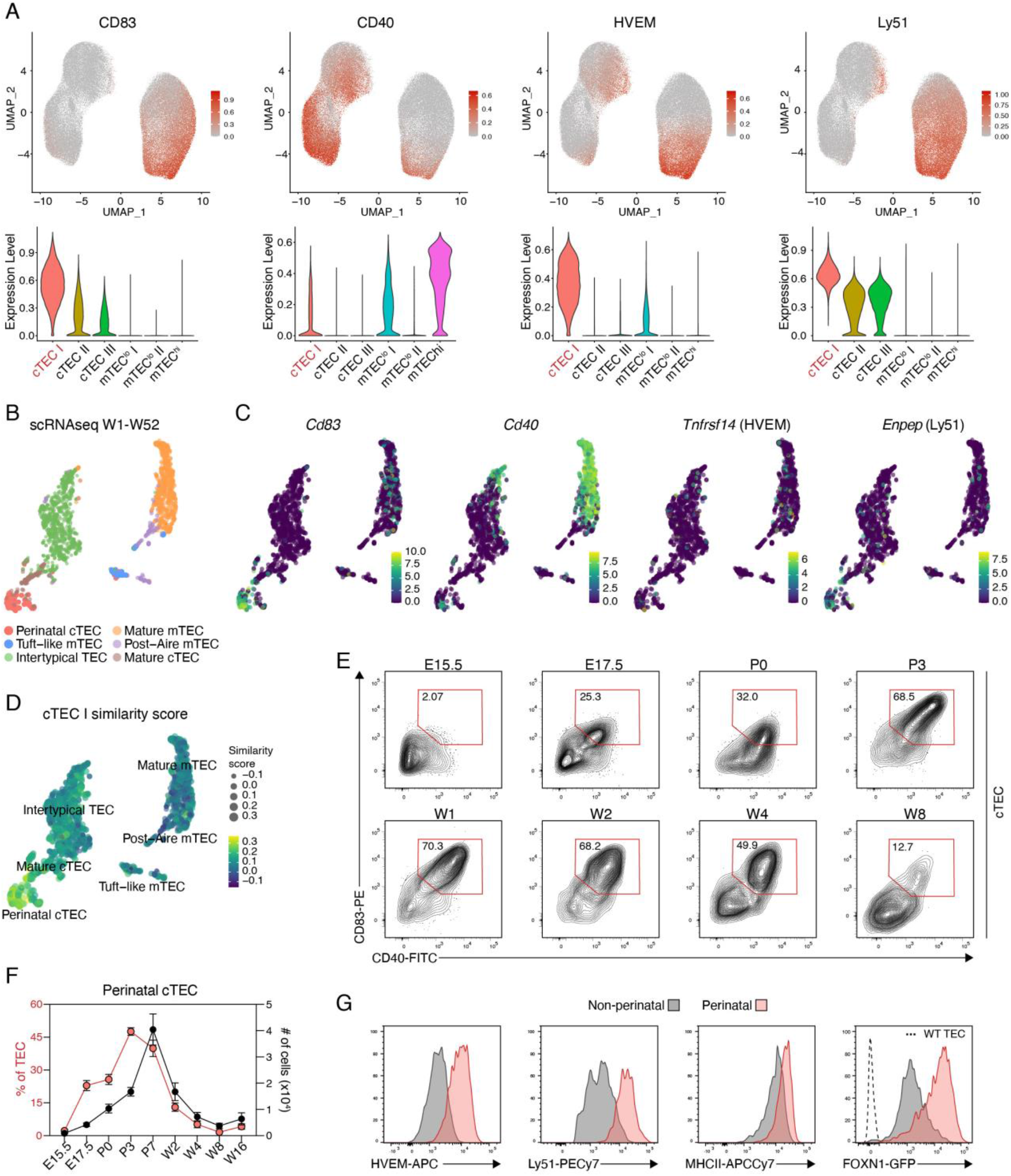
Surface expression profile of perinatal cTEC. **(A)** UMAP graphs (top panels) and violin plots (bottom panels) illustrating the expression of CD83, CD40, HVEM, and Ly51 on TEC from 1-week-old mice. Colour gradient indicates expression levels in the UMAP graphs and colours in the violin plots represent the different clusters, as defined in Figure 1B. **(B)** Hierarchical clustering analysis was performed on single-cell RNA-sequencing data obtained from TEC derived from 1-, 4-, 16-, 32, and 52-week-old mice and projected in a 2-dimensional space using UMAP as explained in the methods. **(C)** UMAP graphs illustrating the scaled expression of *Cd83, Cd40, Tnfrsf14* (HVEM), and *Enpep* (Ly51). Colour gradient indicates expression levels. **(D)** UMAP graph illustrating the similarity score of the cTEC I cluster from the 1-week Infinity Flow dataset to each cell of the scRNAseq reference dataset, based on the surface protein expression levels imputed by Infinity Flow. **(E**,**F)** Abundance of a CD83^+^CD40^+^Sca1^-^ population (hereafter perinatal cTEC) within cTEC was analysed at the indicated timepoints. Shown are (E) representative FACS plots of CD83 and CD40 expression and (F) cumulative data depicting the percent of perinatal cTEC within TEC as well as their total cell numbers. Data are derived from 2-3 independent experiments per timepoint. Error bars indicate standard error of the mean (SEM). E = embryonic day; P = postnatal day; W = postnatal week. **(G)** Representative histograms showing the expression of HVEM, Ly51, MHCII, and *Foxn1*-GFP within perinatal (CD83^+^CD40^+^Sca1^-^) and non-perinatal (CD83^-^CD40^-^) cTEC in 2-week-old mice.

Transcripts for *Cd83, Cd40*, and *Enpep* (encoding Ly51) were detected in perinatal cTEC, albeit at various levels (Figure 2B,C) (Baran-Gale et al., 2020). In contrast, transcripts for *Tnfrsf14*, the gene encoding HVEM, were detected in only a few TEC but across several clusters, thus failing to unequivocally identify perinatal cTEC. This finding highlighted the limitations of gene expression studies to identify surface markers that matched the cells’ RNA profile. To further assess the relationship of the cTEC I subcluster to TEC subtypes identified by scRNAseq, we generated a score of similarity using SingleR which related the RNA expression profile of individual cells to the computed cell surface expression pattern of cTEC I. This analysis demonstrated the highest similarity score to the pairing of cTEC I with perinatal cTEC (Figure 2D).

We then aimed to define surface markers that allow the isolation of perinatal cTEC by flow cytometry. We used the presence of markers highly expressed on cluster cTEC I of 1-week-old mice while excluding markers detected on the majority of mature cTEC isolated from 4- to 16-week-old animals (Figure 1B-D; 2A), as the population of cortical epithelia in older animals only includes perinatal cTEC at a very low frequency. We identified within cTEC a subpopulation of cells that concomitantly expressed CD83 and CD40 but were Sca1 negative early in postnatal life (Figure 2E; S3A). As early as 4 weeks postnatally, TEC with a Sca1^+^ phenotype appeared among CD83^+^CD40^+^ cTEC. These cells were electronically excluded from further analysis as they represent mature cTEC that accumulate with age. The frequency of CD83^+^CD40^+^Sca1^-^ epithelia changed substantially during the life course as the cells’ relative representation progressively increased throughout organogenesis, plateaued in 1-week-old mice (4.6×10^4^ ± 1.4×10^4^ cells) and subsequently decreased, displaying the lowest representation in 8-week-old animals (2.4×10^3^ ± 2.3×10^3^ cells; Figure 2E,F; Figure S3B).

Perinatal cTEC displayed in contrast to other cTEC subpopulations, comprising cTEC clusters II and III, higher cell surface levels for HVEM, Ly51 and MHCII which allowed the identification of these cells in combination with a high cell surface expression of CD83 and CD40, in the absence of Sca1. Perinatal cTEC also showed a higher FOXN1 promoter activity as demonstrated in reporter mice where the expression of GFP is under the transcriptional control of the *Foxn1* locus (Figure 2G).

### Identification of intertypical and tuft-like TEC

The Infinity Flow analysis of adult mice revealed 4 distinct mTEC^lo^ clusters. Clusters I and II were observed in 4- and 16-week-old mice and displayed a similar expression profile for most of the 260 exploratory markers (Figure 1B-D) including a shared expression of Sca1 and CD146 (Figure 3A,B). To match these two phenotypically defined subpopulations to their corresponding transcriptome-determined TEC subtypes, we probed the RNA profiles of single TEC for the expression of *Ly6a/Ly6e* (encoding Sca1) and *Mcam* (encoding CD146). While transcripts for *Ly6a/Ly6e* were detected especially among intertypical TEC (Baran-Gale et al., 2020), *Mcam* - specific RNA was only detected at low levels and in different TEC, but mostly within intertypical TEC subtypes (Figure 3C). This subtype is characterised by transcriptional features characteristic of both cTEC and mTEC – as phenotypically defined by the conventional surface marker Ly51 and UEA-reactivity – contribute to this unique TEC subtype (Baran-Gale et al., 2020). Using again the SingleR package, the similarity scores for both 4- and 16-week-old mTEC^lo^ I and II were calculated to be the highest when matched to the intertypical TEC subtype (Figure 3D; S4A). The mTEC^lo^ subpopulation contained cells that co-expressed Sca1 and CD146 and cells with this phenotype increased with postnatal age (Figure 3E). Although initially only detected among mTEC^lo^ this subpopulation was increasingly also observed within the cTEC compartment of mice older than 4 weeks of age (Figure 3F).

**Figure 3.**
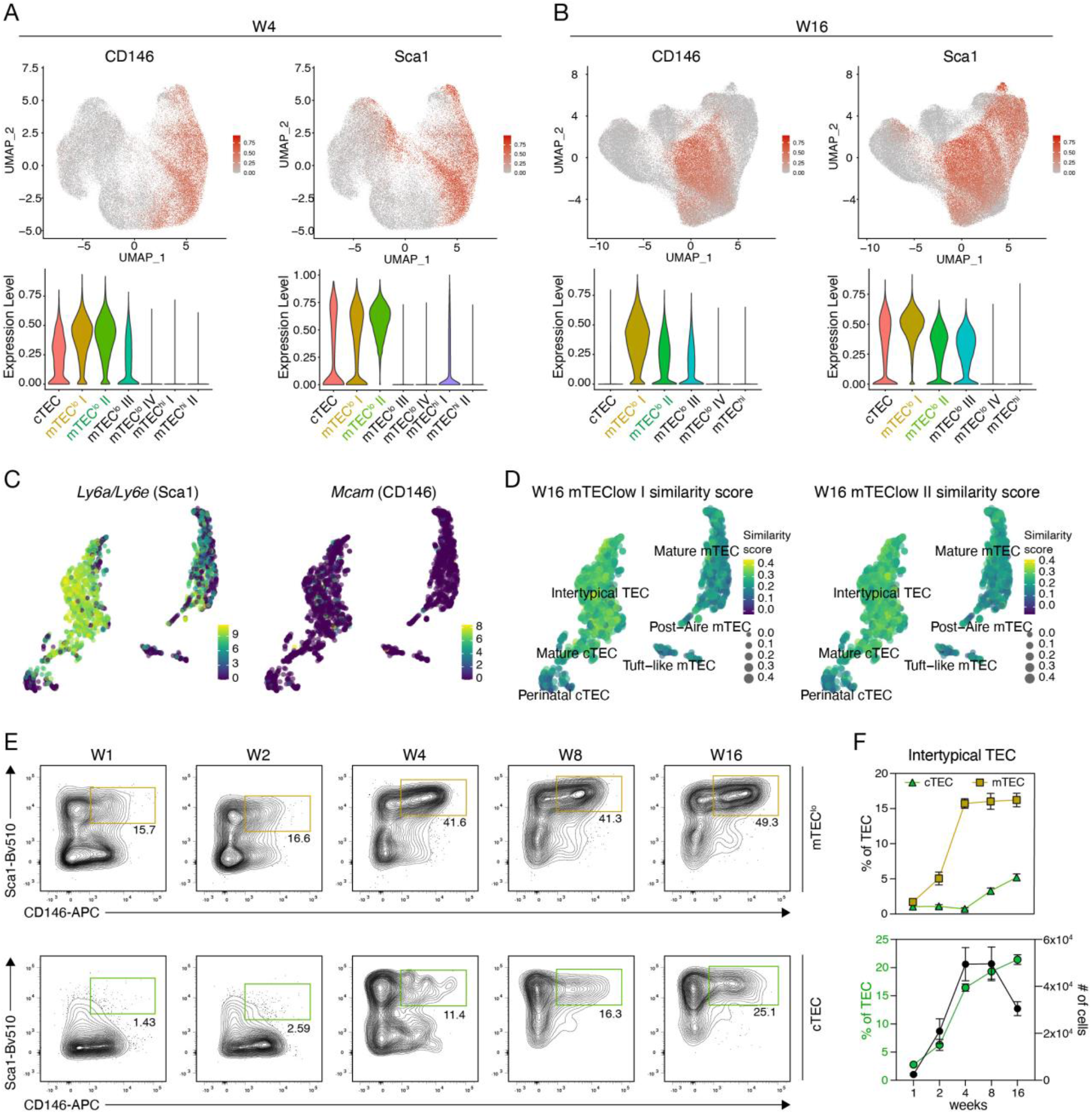
Identification of intertypical TEC within cTEC and mTEC. **(A**,**B)** UMAP graphs (top panels) and violin plots (bottom panels) illustrating the expression of CD146 and Sca1 on TEC from 4-(A) and 16-week-old (B) mice. Colour gradient indicates expression levels in the UMAP graphs and colours in the violin plots represent the different clusters, as defined in Figure 1B. **(C)** UMAP graphs illustrating the scaled expression of *Ly6a/Ly6e* (Sca1) and *Mcam* (CD66a) in the scRNAseq dataset introduced in Figure 2B. Colour gradient indicates expression levels. **(D)** UMAP graph illustrating the similarity score of the mTEC^lo^ I and mTEC^lo^ II clusters from the 16-week Infinity Flow dataset to each cell of the scRNAseq reference dataset, based on the surface protein expression levels imputed by Infinity Flow. **(E**,**F)** Abundance of a Sca1 and CD146 double positive population (hereafter intertypical TEC) within mTEC^lo^ (E; top panels) and within cTEC (E; bottom panels) was analysed at the indicated timepoints. Shown are (E) representative FACS plots and (F) cumulative data for the percent of intertypical TEC within mTEC and cTEC (top panel) and percent of intertypical TEC within TEC as well as their total cell numbers (bottom panel). Data are derived from 2-3 independent experiments per timepoint. Error bars indicate standard error of the mean.

A unique set of surface markers that specifically recognised the mTEC^lo^ cluster III could not be found. However, the simultaneous expression of CD66a and CD117 in the absence of Sca1 and CD63 positivity identified mTEC^lo^ cluster IV. This cluster was only detected in adult mice (Figure 4A; S4B,C), although the defining 4 cell surface markers could also be detected in the mTEC^lo^ cluster II of 1-week-old mice (Figure S4B). It is therefore possible that cluster mTEC^lo^ II of 1-week-old mice represents epithelia that form the separate cluster mTEC^lo^ IV in older animals. The single cell transcriptomic analysis only partially matched the phenotypic analysis of mTEC^lo^ clusters as *Ly6a/Ly6e-* and *Cd63-*specific RNA could indeed be detected in the vast majority of TEC (Figure 3C and 4B) whereas transcripts for *Ceacam1* (encoding CD66a) and *Kit* (encoding CD117) were largely absent in these cells (Figure 4B). The similarity score revealed the best match between mTEC^lo^ cluster IV and post-AIRE and tuft-like mTEC (Figure 4C; 5B,C; S5C).

**Figure 4.**
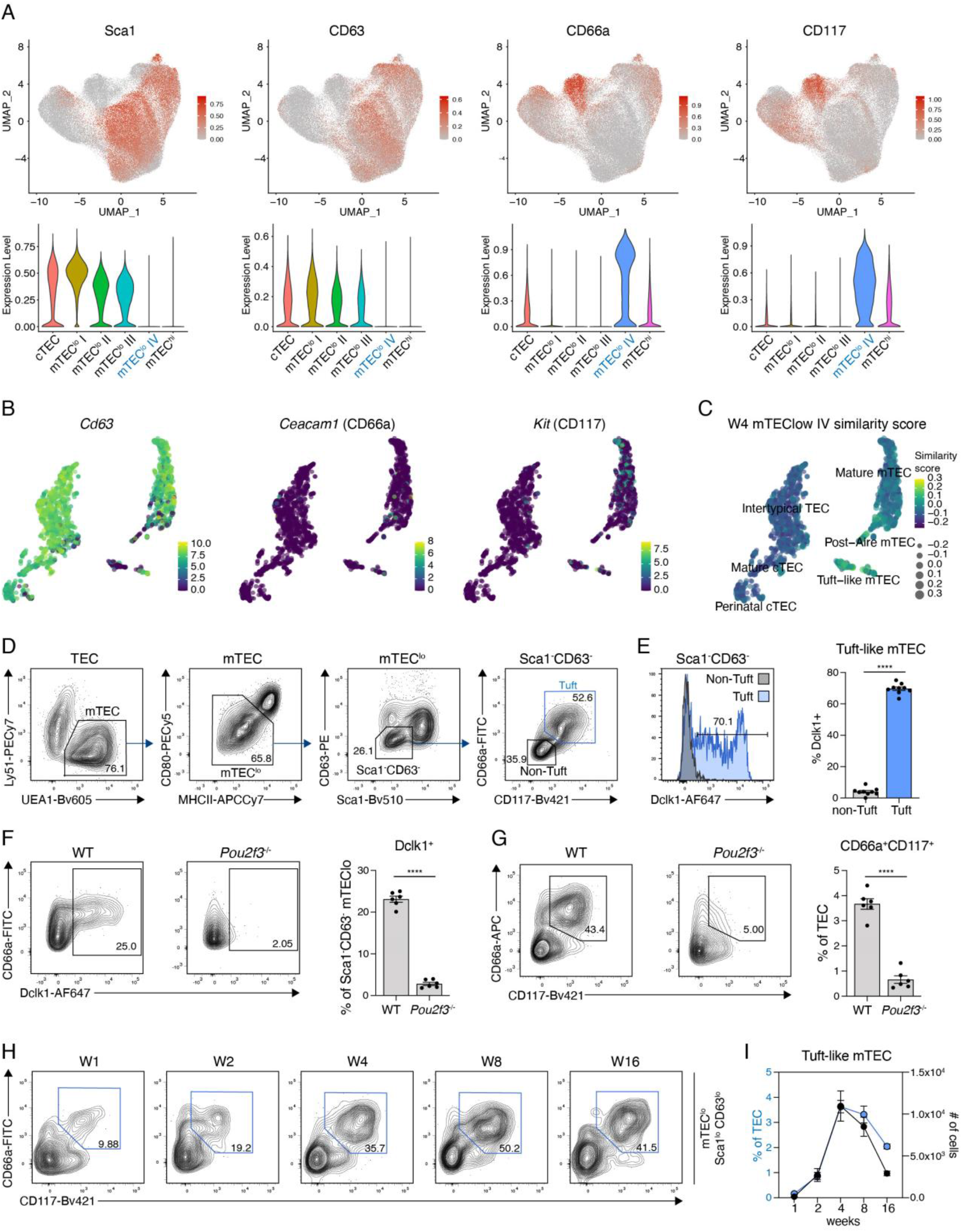
A combination of surface markers to define tuft-like mTEC. **(A)** UMAP graphs (top panels) and violin plots (bottom panels) illustrating the expression of Sca1 CD63, CD66a, and CD117 on TEC from 16-week-old mice. Colour gradient indicates expression levels in the UMAP graphs and colours in the violin plots represent the different clusters, as defined in Figure 1B. **(B)** UMAP graphs illustrating the scaled expression of *CD63* and *Ceacam* (CD66a) and *Kit* (CD117) in the scRNAseq dataset introduced in Figure 2B. Colour gradient indicates expression levels. **(C)** UMAP graph illustrating the similarity score of the mTEC^lo^ IV cluster from the 4-week Infinity Flow dataset to each cell of the scRNAseq reference dataset, based on the surface protein expression levels imputed by Infinity Flow. **(D)** Gating strategy to identify tuft-like mTEC within EpCAM1^+^ cells using Sca1, CD63, CD66a, and CD117. **(E)** Intracellular staining for Dclk1 expression in 4- to 8-week-old mice. Shown are a representative histogram and cumulative data depicting the percent Dclk1^+^ cells within tuft-like mTEC and CD66a^-^CD117^-^ non-tuft cells, as defined in (D). Data are derived from four independent experiments. Error bars indicate SEM. Statistical analysis was done with two-tailed unpaired Student’s t-test. ****, P < 0.0001. **(F**,**G)** *Pou2f3*^*-/-*^ mice were analysed for their abundance of (F) Dclk1^+^ cells and (G) CD66a^+^CD117^+^ cells compared to WT mice. Shown are representative FACS plots (left panels) and cumulative data (right panels). Data are derived from two independent experiments. Error bars indicate SEM. Statistical analysis was done with two-tailed unpaired Student’s t-test. ****, P < 0.0001. **(H**,**I)** Abundance tuft-like mTEC, as defined in (D) within mTEC^lo^ was analysed at the indicated timepoints. Shown are (H) representative FACS plots and (I) cumulative data depicting the percent of tuft-like mTEC within TEC as well as their total cell numbers. Data are derived from 2-3 independent experiments per timepoint. Error bars indicate SEM.

**Figure 5.**
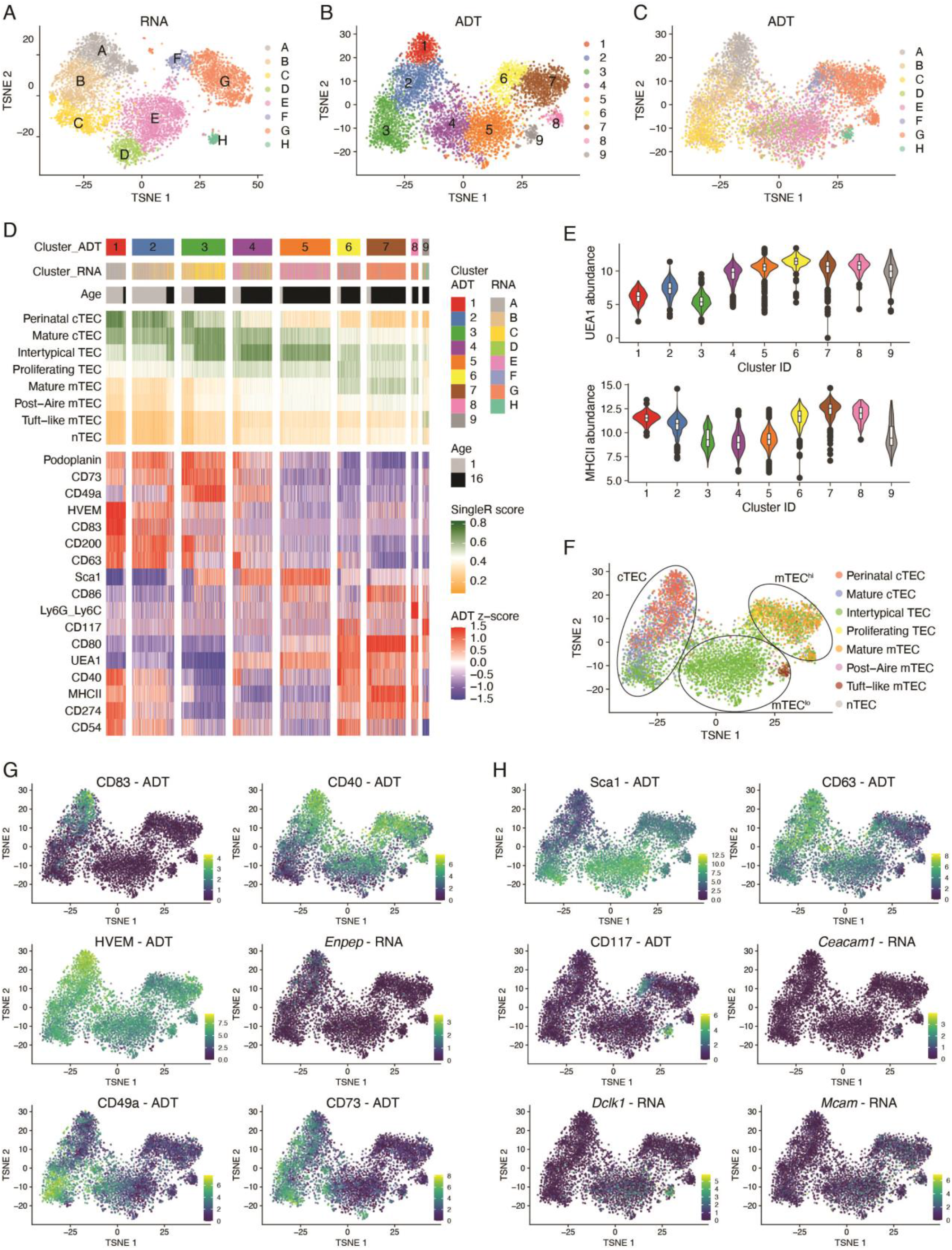
CITEseq validates new TEC markers. CD45^-^Ter119^-^ thymic stromal cells isolated from 1- and 16-week-old WT mice were used for scRNAseq in combination with CITEseq as described in the methods. Cells belonging to clusters assigned as epithelial cells were selected for further analysis. **(A-C)** Hierarchical clustering analysis was performed on 5834 TEC either using (A) the gene expression analysis or (B) only considering the detection of ADTs. Results were projected in a 2D space using t-distributed stochastic neighbour embedding (t-SNE). Each colour represents a specific cluster. In (C) t-SNE distribution of the ADT clustering is shown using the cluster colouring of the RNA analysis. **(D)** Compiled data showing the cluster distributions, defined as in A and B, in relation to the derivation of the cells from 1- or 16-week-old mice, and to the similarity score to the reference TEC scRNAseq dataset from Bara-Gale et al (Baran-Gale et al., 2020). The expression of CITEseq markers is centred to the mean and scaled to the range of expression values. **(E)** Violin plots depicting the abundance of UEA1 and MHCII ADTs across ADT clusters. **(F)** Cells were annotated based on transcriptional similarity to the scRNAseq dataset from Baran-Gale et al (Baran-Gale et al., 2020). Each colour represents a specific TEC subset as defined in the reference dataset. **(G**,**H)** T-SNE plots illustrating the scaled expression of (G) perinatal cTEC markers such as CD83, CD40, HVEM, *Enpep*, CD49a, and CD73, and of (H) tuft-like and intertypical TEC markers such as Sca1, CD63, CD117, *Ceacam1, Dclk1*, and *Mcam* across ADT clusters.

We next sought to define a phenotypic profile of cells belonging to cluster mTEC^lo^ IV that would allow their physical isolation by flow cytometry. For this purpose, we identified within the Sca1^-^CD63^-^ mTEC^lo^ a subpopulation of cells that stained positively for both CD66a and CD117, thus mirroring the features identified for mTEC^lo^ cluster IV (Figure 4D). Most of the Sca1^-^CD63^-^ CD66a^+^CD117^+^ mTEC^lo^ cells (∼70%) also expressed the serine/threonine-protein kinase Dclk1 (Figure 4E) which was previously identified as a typical intracellular marker for tuft-like mTEC (Bornstein et al., 2018; Lucas et al., 2020). Conversely, the vast majority of Dclk1-positive TEC were detected among Sca1^-^CD63^-^CD66a^+^CD117^+^ mTEC^lo^ (Figure S4E). In the absence of the transcription factor Pou2f3 Dclk1 expression was absent and Sca1^-^CD63^-^CD66a^+^CD117^+^ mTEC^lo^ cells were not generated (Figure 4F,G) thus confirming their identity as tuft-like thymic epithelia (Lucas et al., 2020; Miller et al., 2018). We also observed that an absence of Dclk1 in wild-type Sca1^-^ CD63^-^CD66a^+^CD117^+^ mTEC^lo^ correlated with a lower surface expression of CD66a and CD117, indicating that they are not yet fully differentiated into tuft-like mTEC (Figure S4F). Using these phenotypic features, we noted the presence of tuft-like mTEC to change over time with the highest frequency and cellularity in 4-week-old mice (Figure 4H,I).

Tuft -like mTEC originate from mTEC that have once expressed the tissue restricted antigen, Csnb (Bornstein et al., 2018). We utilised *Csnb*^Cre^::Rosa26^LSL-YFP^ reporter mice (Bornstein et al., 2018) that allow *in vivo* fate mapping within the mTEC linage to test whether Sca1^-^CD63^-^ CD66a^+^CD117^+^ mTEC^lo^ originate from a *Csnb* expressing precursor. In keeping with the previous study, we identified 70-80% of the Sca1^-^CD63^-^CD66a^+^CD117^+^ mTEC^lo^ mTEC to be YFP labelled (Figure S5A), suggesting that they represent bona-fide tuft-like mTEC. The mTEC^lo^ compartment is composed of medullary epithelia that either have not yet expressed the transcriptional facilitator AIRE or, alternatively, belong to a group of cells that have differentiated from AIRE-positive, mature mTEC. Tuft-like mTEC have previously been shown to derive, at least in part, from AIRE-positive precursors (Miller et al., 2018) and be enriched for expression of the surface glycoprotein Tspan8, an AIRE-enhanced tissue-restricted antigen (Dhalla et al., 2020; Rattay et al., 2016). We therefore tested whether Sca1^-^CD63^-^CD66a^+^CD117^+^ mTEC^lo^ (i.e. tuft-like) mTEC are positive for the expression of Tspan8. As many as 60% of tuft-like mTEC expressed Tspan8, further validating their identity and identifying these cells to contain post-AIRE mTEC (Figure S5B). The combined expression of YFP and Tspan8 was only detected in a small fraction of Sca1^+^CD146^+^ mTEC^lo^ intertypical TEC (Figure S5), suggesting they belong to a TEC developmental stage that does not yet promiscuously express tissue specific antigens.

### CITEseq validates novel TEC markers

We next analysed both surface protein and mRNA expression of individual thymic stromal cells using Cellular Indexing of Transcriptomes and Epitopes by sequencing (CITEseq) (Stoeckius et al., 2017). This approach was taken to confirm unequivocally that the combined use of the newly described cell surface markers indeed identified a specific TEC subtype. For this purpose, CD45^-^ Ter119^-^ cells were isolated from thymi of 1- and 16-week-old mice and stained with oligonucleotide-coupled antibodies each directed against either EpCAM1, MHCII, UEA1, CD80, CD86, CD40, CD83, HVEM, CD73, Sca1, CD63, CD117, CD200, CD54, CD49a, CD274, Ly6G/Ly6C, Podoplanin, or CD31. Labelled TEC were then analysed by scRNAseq. The hierarchical clustering analysis identified 12 clearly separated clusters whether data from single cell gene expression profiling or, alternatively, from antibody derived tags (ADT) was computed and displayed by means of t-distributed stochastic neighbour embedding (tSNE) (Figure S6A-C). The cell-type annotation was based on the gene expression profiles derived from the Immunological Genome Project (ImmGen) and confirmed the identity of individual clusters as endothelial cells, fibroblasts, stromal cells, and epithelial cells, respectively. Hence, the chosen combination of selected surface markers was sufficient to identify individual stromal cell types (Figure S6D,E).

Analysing the captured gene expression profiles of only TEC identified 8 separate clusters (A-H), whereas examining the ADT data recognised 9 (1-9) clusters (Figure 5A,B). The two clustering approaches largely overlapped, suggesting a robust separation accomplished between individual clusters (Figure 5C). Clusters defined by mRNA-profiling related to an individual ADT-defined population with a few exceptions: clusters D and E represented a mixture of clusters 4 and 5, and cluster G split into clusters 7 and 8 which differed in the expression of Ly6C/Ly6G but not the cells’ RNA expression profiles (Figure 5D). The limited number of antibodies used in the CITEseq analysis identified three cTEC (defined as UEA1 non-reactive cells), cluster 1 corresponding to cluster A [1/A], 2/B, and 3/C), three mTEC^lo^ (MHCII^lo^CD80^lo^: 4/D, 5/E, and 9/H) and three mTEC^hi^ subpopulations (MHCII^hi^CD80^hi^: 6/F, 7/G, and 8) (Figure 5D-F and S6F).

We next assigned CITEseq-defined TEC cluster identities to those subtypes we have previously classified (Baran-Gale et al., 2020). Clusters 1/A and, to a lesser extent, 2/B corresponded to perinatal cTEC. Cluster 2/B was further related to mature cTEC and cluster 3/C to mature cTEC and intertypical TEC (Figure 5D,F). Clusters 4/D and 5/E related to intertypical TEC while cluster 6/F was most similar to both mature mTEC and proliferating TEC. Clusters 7/G and 8/G displayed a high similarity to mature mTEC (Figure S6F) whereas cluster 9/H linked to tuft-like mTEC. Notably, the transcriptional signatures characteristic of post-Aire mTEC and neural (n) TEC could not be detected. Replicating the changes of specific TEC subtypes with age, clusters 1/A and 2/B were more abundant in 1-week-old mice whereas the frequencies of clusters 3/C, 4/D, 5/E, 7/G, and 9/H were increased in 16-week-old animals (Figure 5D; Figure S6G).

TEC in cluster 1/A displayed the highest CD83, CD40, and HVEM protein expression among CITEseq defined clusters, thus confirming the cells’ identity as perinatal cTEC. The expression of these markers was reduced in cluster 2/B and completely absent in cluster 3/C, suggesting the former to represent a developmentally intermediate cell state between perinatal and mature/intertypical-like cTEC (Figure 5D,G). Furthermore, cTEC maturation was paralleled by a decrease in *Foxn1* transcription and an increase in CD73, CD49a, and Sca1 protein expression (Figure S6H).

The differential CD117, CD63, and Sca1 protein expression (as measured by ADT) identified cluster 9/H as tuft-like mTEC (CD117^+^CD63^-^Sca1^-^; see above and Figure 5D,H) and thus confirmed the flow cytometric definition and gating strategy used to identify these cells as both accurate and practical (Figure 4G-I). This conclusion was further corroborated by the detection of *Dclk1* and *Ceacam1* transcripts in cluster H (Figure 5H) and a high similarity score with the tuft-like mTEC subtype (Figure 5D,F)(Baran-Gale et al., 2020).

ADT-based detection of Sca1 protein expression matched to cells with a transcriptional signature of intertypical TEC within the cTEC (cluster C) and mTEC^lo^ subpopulations (cluster E; Figure 5D,F,H). Hence, intertypical TEC could unequivocally be identified by Sca1 expression alone. As the transcriptional signature identifying intertypical TEC was spread across three CITEseq-defined clusters (3/C, 4/D and 5/E; Figure 5D,F), the detection of CD146 expression appeared to deconvolute TEC heterogeneity further since fractions of Sca1^+^ cTEC and Sca1^+^ mTEC^lo^ stained positively for CD146^+^ (Figure 3E).

The ADT-based documentation of surface markers identified individual TEC subtypes. However, the corresponding gene expression profiles were on their own insufficient to recognize these cells, not least because of the occasional discrepancy between surface protein and RNA expression (Figure S6H,I). CITEseq could therefore validate the utility of the selected, novel surface markers and the gating strategy chosen. Together they identified 4 TEC subpopulations that correspond to a specific transcriptionally-defined cluster and 2 subpopulations that represent a mixture of 2 related clusters, namely UEA1^-^CD83^+^CD40^+^Sca1^-^ perinatal cTEC (cluster 1/A), UEA1^-^ CD83^-^CD40^-^Sca1^-^ mature cTEC (2/B), UEA1^-^CD83^-^CD40^-^Sca1^+^ intertypical-like cTEC (3/C), UEA1^+^MHCII^lo^CD80^lo^Sca1^+^ intertypical mTEC (4+5/D+E), UEA1^+^MHCII^hi^CD80^hi^ mature mTEC (6+7+8/F+G), and UEA1^+^MHCII^lo^CD80^lo^Sca1^-^CD63^-^CD66a^+^CD117^+^ tuft-like TEC (9/H) (Figure S7).

### Perinatal cTEC present an enhanced potential for positive selection

We next sought to localize perinatal cTEC within the thymus stromal architecture. Because Ly51 expression was higher on perinatal cTEC in comparison to other cortical epithelial populations (Figure 2G), we used this differential to localize perinatal cTEC on thymus tissue sections (Figure 6A). Quantification of the Ly51 signal intensity in immunohistology detected these cells in close proximity to the medulla with gradual increase of Ly51 signal but invariable cytokeratin 8 (K8) staining across the cortex from subcapsular region to the inner and eventually deep cortex (Figure 6A,B). Contrary to flow cytometry, the immunostaining patterns of antibodies directed against CD83, CD40, and HVEM were not informative (data not shown).

**Figure 6.**
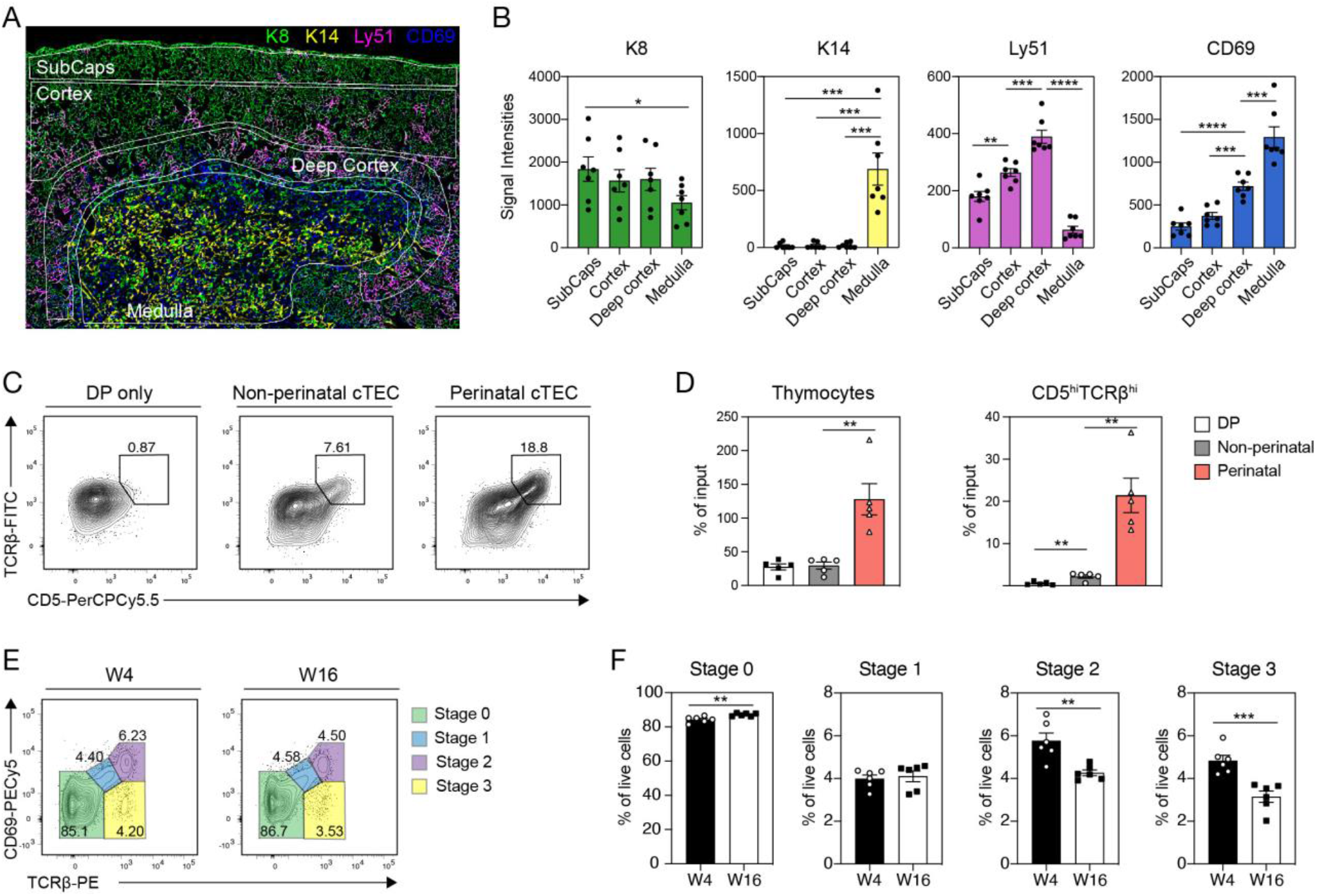
Perinatal cTEC display an increased capacity for positive selection. **(A**,**B)** Immunofluorescent analysis of frozen thymic tissue sections from 4-week-old mice stained with antibodies directed against K8 (green), K14 (yellow), Ly51 (magenta), and CD69 (blue). Shown are (A) an image of a representative region (n=7) and (B) cumulative data depicting the signal intensities detected across the subcapsular region (SubCaps), the inner cortex, the deep cortex and the medulla. Data are derived from three biological samples. Error bars indicate SEM. Statistical analysis was done with two-tailed unpaired Student’s t-test. **(C**,**D)** Reaggregate thymic organ cultures (RTOC) of non-perinatal (CD83^-^CD40^-^) and perinatal (CD83^+^CD40^+^Sca1^-^) cTEC with CD69^-^ DP thymocytes were performed. Shown are representative FACS plots illustrating the expression of (C) TCRβ and CD5 after two days of culture for DP only, non-perinatal cTEC and perinatal cTEC cultures and the number of (E) thymocytes and CD5^hi^TCRβ^hi^ cells acquired. Data are derived from three independent experiments. Statistical analysis was done with two-tailed unpaired Student’s t-test. **, P < 0.01. **(E**,**F)** Abundance of developmental thymocyte stages based on the expression of TCRβ and CD69 was analysed in 4- and 16-week-old mice. Shown are (E) representative FACS plots and (F) cumulative data revealing the percent of cells of thymocyte stages 0-3. Data are derived from two independent experiments (n=6). Error bars indicate SEM. Statistical analysis was done with two-tailed unpaired Student’s t-test. **, P < 0.01; ***, P < 0.001.

The new surface markers enabled the isolation and functional testing *ex vivo* of individual cTEC populations. We therefore investigated the capacity of perinatal (CD83^+^CD40^+^Sca1^-^) and non-perinatal (CD83^-^CD40^-^) cTEC to effect positive thymocyte selection. We co-cultured these cells as reaggregate thymic organ cultures (RTOCs) together with CD69^-^CD4^+^CD8^+^ (i.e. pre-selection double-positive) thymocytes for two days before thymocytes were monitored for phenotypic features associated with positive thymic selection, i.e. the upregulation of TCR and CD5 (Figure 6C). The number of total thymocytes and those with a CD5^hi^TCRβ^hi^ phenotype were significantly increased in RTOCs composed of perinatal cTEC when compared to aggregates composed of other cortical epithelia (Figure 6D). Taken together, these results identified perinatal cTEC to be juxtaposed to the medulla and particularly efficient in effecting positive thymocyte selection.

The number of perinatal cTEC significantly decreases with age (Baran-Gale et al., 2020). We therefore explored whether this variation was paralleled by a change in the efficiency to impose positive thymocyte selection. We thus monitored and compared thymocyte maturation in 4- and 16-week-old thymi and classified their sequential maturational stages according to the cells’ expression of TCRβ and CD69 (i.e. stage 0: TCRβ^-^CD69^-^ → stage 1:,TCRβ^+^CD69^+/-^ → stage 2: TCRβ^+^CD69^+^ → stage 3: TCRβ^+^CD69^-^). The frequency of pre-selection thymocytes (i.e. stage 0) was increased whereas the relative abundance of cells with a post-selection phenotype (stage 2 and 3) was significantly reduced in older animals (Figure 6E,F). These *in vivo* results indicated a compromised capacity of older mice to positively selected thymocytes, which correlated with a decrease in the availability of perinatal cTEC.

### Crosstalk with thymocytes induces maturation of perinatal cTEC

We finally investigated whether thymic crosstalk (Abramson and Anderson, 2017; Hollander et al., 1995) could explain the inverse correlation between the decreased frequency of perinatal cTEC with age and the expansion of thymocytes after birth. We therefore first determined the frequency of perinatal cTEC in Rag2-deficient (Rag2^-/-^) mice, which have a hypoplastic thymus secondary to a thymocyte developmental arrest at the DN3a stage. We found a high fraction of perinatal cTEC in these mice that was not influenced by age (Figure 7A). Hence, thymocytes at developmental stages up to the beta-checkpoint did not influence the age-related changes in perinatal cTEC frequencies.

**Figure 7.**
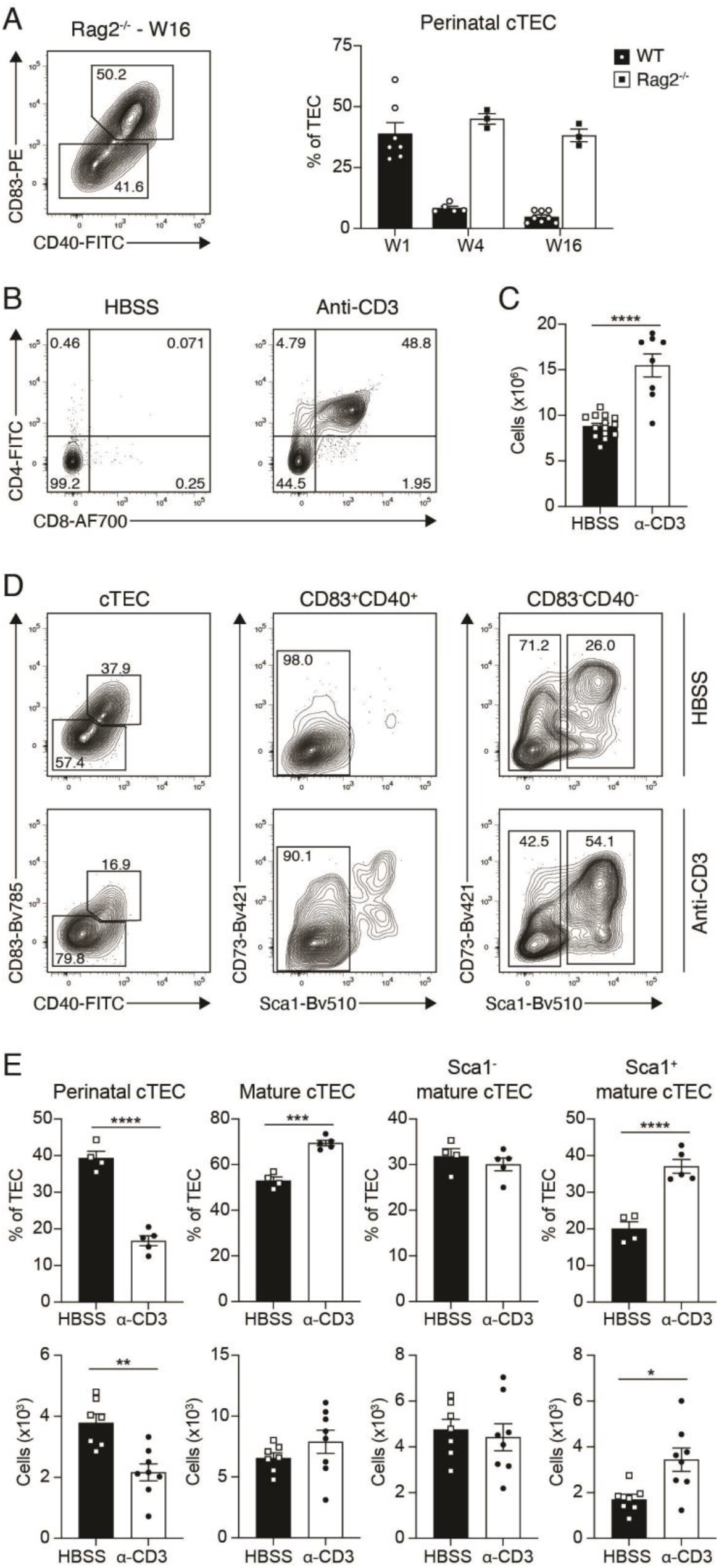
Crosstalk with thymocytes facilitates cTEC maturation. **(A)** Rag2^-/-^ mice were analysed for the abundance of perinatal cTEC at 4- and 16-weeks and compared to perinatal cTEC in 1-, 4-, and 16-week-old WT mice. Shown are a representative FACS plots and cumulative data. Data shown are derived from one out of two independent experiments. Error bars indicate SEM. **(B**,**C)** Rag2^-/-^ mice were injected with HBSS or α-CD3 antibodies (clone KT3) and analysed four weeks later for the development of double positive thymocytes. Shown are representative FACS plots depicting the emergence of CD4^+^CD8^+^ cells and cumulative data for the total number of cells per thymus. Data are derived from two independent experiments. Error bars indicate SEM. **(D**,**E)** The cTEC compartment was analysed for changes in the abundance of subpopulations following α-CD3 treatment. Shown are (D) representative FACS plots and (E) cumulative data as percentage of TEC and as total cell numbers. Data are derived from two independent experiments. Error bars indicate SEM. Statistical analysis was done with two-tailed unpaired Student’s t-test. *, P < 0.05; **, P < 0.01; ***, P < 0.001; ****, P < 0.0001.

To probe whether thymocytes at later developmental stages, especially unselected CD4^+^CD8^+^ (double positive, DP) thymocytes, controlled the frequency of perinatal TEC, Rag2^-/-^ mice were injected with antibodies directed against CD3ε. This treatment results in a substantial increase in pre-selection DP thymocytes (Jacobs et al., 1994; Levelt et al., 1995; Shinkai and Alt, 1994). Four weeks after antibody or control injections, the thymus of actively treated Rag2^-/-^ mice contained an abundance of DP thymocytes which correlated with numerical and phenotypic changes in the cTEC compartment (Figure 7B-E). The latter were marked by a reduction in perinatal cTEC, parallel to an increase in mature cTEC, specifically Sca1^+^ cells (Figure 7D,E) which corresponded to intertypical TEC according to our CITEseq data (Figure 5D,F,H). Taken together, these results identified the abundance of and/or signalling by pre-selection DP thymocytes as the mechanism by which the frequency of perinatal cTEC was controlled.

## Discussion

Single cell transcriptomic analyses have uncovered an unexpected heterogeneity within many cell populations of a seemingly identical phenotype. Cells of the thymic stromal compartment constitute no exception to this observation ((Baran-Gale et al., 2020; Bornstein et al., 2018; Dhalla et al., 2020; Handel et al., 2022; Miller et al., 2018). The apparent lack of suitable cell surface markers identifying unequivocally TEC subpopulations identical to individual TEC subtypes precludes the isolation of live TEC and their *ex vivo* functional analysis. Here we report that this limitation has been substantially overcome. We describe novel cell surface markers that identify the comparable subtype of scRNAseq-defined perinatal, mature and intertypical cTEC, and mature, intertypical and tuft-like mTEC.

Predicting the cell surface phenotypes from corresponding scRNAseq profiles is challenging as technical limitations detecting low transcript copy numbers and the acknowledged disparity between transcript detection and protein expression render this attempt difficult. For example, a comparison of 7 scRNAseq methods uncovered that high-throughput methods, including the widely used 10x Chromium, have lower sensitivities in comparison to the low-throughput methods Smart-seq2 and CEL-Seq2 when capturing rare transcripts (Ding et al., 2020; Zheng et al., 2017). We therefore opted for an alternative method and stained TEC for the expression of hundreds of cell surface markers. This screening approach of massive parallel flow cytometry combined with Inifnity Flow analysis discovered surface markers previously not inferred to be expressed by TEC. CITEseq which combines the detection of promising candidate markers and single cell transcriptomic profiles finally established the accuracy of the cell surface markers chosen to identify TEC subtypes.

scRNAseq across the life-trajectory identifies a substantial heterogeneity among TEC and with it a dynamic change of the cells’ relative frequencies (Baran-Gale et al., 2020). For example, the cTEC compartment is composed of at least two main subtypes, designated perinatal and mature cTEC (Baran-Gale et al., 2020). Perinatal cTEC represent a major subpopulation early in life (∼40% of all TEC the first week after birth) but their relative frequency rapidly decreases thereafter with only a small fraction of these cells being detected in adult animals. Conversely, mature cTEC increase in frequency and represent the majority of cortical epithelia from 4 weeks of life onwards (Baran-Gale et al., 2020). Intertypical TEC are characterised by a gene expression profile that includes signatures typical for both cortical and medullary thymic epithelial lineages. They also express genes including *Pdpn, Ccl21a, Ly6a*, and *Plet1* that have previously been associated with mTEC thought to have a progenitor potential and localised at the cortico-medullary junction (CMJ) (Mayer et al., 2016; Michel et al., 2017; Nusser et al., 2022; Onder et al., 2015; Ulyanchenko et al., 2016). The hitherto absence of suitable and informative cell surface markers to physically isolate most of the TEC subtypes for *in vitro* analyses and *in vivo* transfer studies has disallowed to date further functional characterizations of these cells and the physical establishment of direct precursor::progeny relationships.

By applying the newly identified surface markers we show that perinatal cTEC are enriched within the TEC scaffold towards the cortico-medullary junction (CMJ) and particularly efficient in positively selecting maturing thymocytes. As expected for their role in shaping the TCR repertoire, we find perinatal cTEC typically juxtaposed to thymocytes with an activated phenotype (i.e. CD69^+^). The age-dependent decline in the frequency of perinatal cTEC is noted both when using flow cytometry and scRNAseq to classify these cells. The actual pace by which this regression is observed differs, however, between these two methods, demonstrating a seemingly faster kinetic for perinatal cTEC identified by their RNA expression profile (Baran-Gale et al., 2020). This may be explained by differences in the half-lives of specific transcripts and their corresponding proteins. Nonetheless, a post-natal decrease of these cells to an almost complete absence early in adulthood is expected to compromise the robustness of thymopoiesis and possibly the efficiency by which thymocytes are positively selected. The observed decrease in perinatal cTEC correlates with other compositional changes within the epithelial scaffold and may constitute an intrinsic driver for thymus senescence. This understanding is consistent with scRNAseq data that shows the quiescence of a population of medullary precursor cells and correlates this alteration with an impaired maintenance of the medullary TEC compartment (Baran-Gale et al., 2020). In parallel, these age-related changes link to less efficient T cell selection, a decreased self-antigen representation, an increased T cell receptor repertoire diversity, and a reduced frequency of thymus-resident naïve T cells (this report and (Baran-Gale et al., 2020)).

Intertypical TEC express the glycosyl phosphatidylinositol-anchored cell surface protein Sca1 independently whether they are positive for the cortical marker Ly51^+^ or reactive with the lectin UEA-1, a general feature of medullary TEC. Here, we now show that intertypical TEC can be further split into two sizable subpopulations based on their expression of the cell adhesion molecule CD146. Because oligonucleotide labelled anti-CD146 antibodies were not available for the CITEseq analysis and transcripts for this marker are typically lowly expressed among intertypical TEC, a distinction between CD146 negative and positive cells was not possible when analysing scRNAseq data. Intertypical TEC may contain progenitors with a developmental bias towards the mTEC lineage (Baran-Gale et al., 2020). Indeed, a recent report provides further support of this contention since the gene expression profile of intertypical TEC largely overlaps with a heterogeneous progenitor population which has been claimed to act as mTEC biased postnatal TEC progenitor (Nusser et al., 2022). Our profiling of Tspan8 expression and lineage tracing furthermore suggest that the majority of Sca1^+^CD146^+^ intertypical TEC relate to immature mTEC that have not yet fully differentiated to express collectively a broad range of tissue restricted antigens.

We further specify the cell surface phenotype for tuft-like mTEC (L1CAM^+^CD104^+^ (Bornstein et al., 2018)). These cells share transcriptional (e.g. expression of IL25, Trmp5, Dclk1, and IL17RB) and morphological characteristics with gut epithelial tuft cells (Baran-Gale et al., 2020; Bornstein et al., 2018; Miller et al., 2018), play a function in central T cell tolerance induction (Miller et al., 2018) and control both the homeostasis of type 2 innate lymphoid cells and the generation of type 2 natural killer T cells (Bornstein et al., 2018; Lucas et al., 2020; Miller et al., 2018). The lineage tracing of these cells further shows that the majority of tuft-like mTEC derive from AIRE expressing TEC, a finding in keeping with previously reported observations (Bornstein et al., 2018; Miller et al., 2018). However, it remains an open question whether all tuft-like mTEC differentiate from mature mTEC because the labelling method to draw this conclusion (i.e. inducible, *Aire* dependent tracing) is not necessarily completely effective (Miller et al., 2018). Interestingly, the *Csnb* lineage tracing identified an increased frequency of labelled tuft-like cells in comparison to mature mTEC where labelling is initiated. While we have no unequivocal explanation for this increase, we nevertheless conclude that the majority of tuft-like mTEC (at least 60%) are the progeny of mature medullary epithelia. There is however room to speculate that all tuft-like mTEC may be derived in this way, since any contributions from another *Csnb* non expressing (i.e. non-labelled) precursor would dilute the frequency of labelled tuft-like cells, a result that we did not observe.

To probe the utility of the new set of cell surface markers to phenotype altered TEC scaffolds, we next analysed the composition of the thymic epithelia in FOXN1^Δ505/WT^ mice which express a dominant negative mutation of FOXN1 and consequently show substantial defects in TEC differentiation (Rota et al., 2021). A previous scRNAseq-based analysis of these animals revealed a relative enrichment of perinatal and mature cTEC against a reduction of tuft-like mTEC whereas the frequency of intertypical TEC remained unchanged. The flow cytometric analysis of the TEC scaffold in FOXN1^Δ505/WT^ mice identifies the same variations and thus maps accurately to the transcriptional analysis of these cells, thus demonstrating that the phenotypic and gene expression-based analyses draw comparable conclusions (Figure S8).

With the approach taken, five of the previously defined nine TEC subtypes (Baran-Gale et al., 2020) can now be unequivocally identified using cell surface markers. The still small number of discriminatory cell surface markers so far identified likely accounts for this minor limitation. Implementing more markers in the screening process may identify additional cell surface markers that will identify the remaining TEC subtypes for which we have not yet identified an unambiguous cell surface marker profile. Alternatively, the use of intracellular markers may be informative in identifying the remaining TEC subtypes, namely proliferating TEC, post-AIRE mTEC, and nTEC. However, an obvious drawback for this approach will be that TEC identified in this fashion will be non-viable and can therefore not be used for functional studies *in vitro* or after transfer *in vivo*.

It is important to establish precursor-progeny relationships for specific TEC subtypes now that we can identify and purify specific subpopulations. For example, the potential can be tested whether CD146^+^ intertypical TEC give rise to mature mTEC, competent to effect negative selection. Another effort could be directed in dissecting the molecular requirements of tuft-like mTEC controlling the development of type 2 lymphoid cells employing *ex vivo* functional assays. Finally, the screening workflow described here will also be valuable in identifying novel biomarkers apt to monitor changes in cell subpopulations and their functions resulting from spontaneous or engineered changes in gene function.

Taken together, we have identified novel surface markers that enable the isolation and functional assessment of novel TEC subpopulations that correspond to previously identified subtypes so far only defined by their transcriptome. This is accomplished by combining a high throughput screening workflow with a computational expression projection followed by unsupervised clustering.

## Methods

### Mice

Animals were maintained under specific pathogen–free conditions at the University of Oxford Biomedical Science facilities. Experiments were performed according to institutional and U.K. Home Office regulations and age- and gender-matched wild-type mice were used in all experiments as a reference for genetically modified animals.

*Csnb*^Cre^ mice (Bornstein et al., 2018) were crossed to the Rosa26^YFP^ mouse line (Srinivas et al., 2001) to induce lineage tracing in the mature mTEC compartment.

Mice heterozygous for a *Foxn1* allele with a single nucleotide loss at position 1470 (designated FOXN1^Δ505/WT^) were generated at the Genome Engineering Facility of the MRC Weatherall Institute of Molecular Medicine, University of Oxford as previously described (Rota et al., 2021).

Rag2^-/-^ mice were bred and maintained in the mouse facility of the Department of Biomedicine at the University of Basel in accordance with permissions and regulations of the Cantonal Veterinary Office of Basel-Stadt.

For timed pregnancies 7-14 week old mice were mated over-night and separated early next morning. For pregnant females the mating was considered E0.5 that morning.

### TEC isolation

Isolated thymi were cleaned from adipose tissue, separated into the two lobes, and subsequently subjected to three rounds of enzymatic digestion with Liberase (2.5 mg/ml, Roche) and DNaseI (10 mg/ml, Roche) diluted in PBS (Sigma) at 37°C. After filtration through a 100 μm cell strainer and resuspension in FACS buffer (PBS supplemented with 2% FBS (Sigma)), cell number was determined using a CASY cell counter (Innovatis). For most analyses CD45^+^ hematopoietic cells were depleted by incubation with anti-CD45 beads (Miltenyi) as per manufacturer’s recommendations and subsequently subjected to the AutoMACS separator (Miltenyi) “depleteS” program.

### Flow cytometry

Cells were counted and stained in FACS buffer containing antibodies of interest (Table S1) for 30 min at 4°C in the dark. For the identification of dead cells an additional staining with propidium iodide (PI, Sigma) or Zombie red (Biolegend) was used. For intra-cellular staining, cells were fixed and permeabilised after cell-surface staining using the Cytofix/Cytoperm (BD Biosciences) or the Fix/Perm buffer set (Invitrogen) according to the manufacturer’s protocol. Cells were analysed and sorted on a BD FACSAria III instrument (BD Biosciences). Cells were sorted into FACS buffer. Cell purities of at least 95% were confirmed by post-sort analysis.

### Massively parallel flow cytometry

Cells were isolated and CD45 depletion plus backbone staining were performed as described. The surface backbone panel included antibodies directed against CD45, EpCAM1, Ly51, MHCII, CD40, CD80, CD86, Sca1, Podoplanin, CD31, the *Ulex europaeus* agglutinin I (UEA1) lectin was used labeled with biotin, followed by secondary streptavidin-Bv421 staining and Zombie red staining. Subsequently, the stained cells were distributed across the three 96-well plates provided with the LEGENDScreen kit (Biolegend), each well containing a unique PE-labeled exploratory antibody as well as isotype controls and blanks. PE-labeled antibodies targeting GP2, Tspan8, CD177 and F3 were used as additional exploratory surface antibodies. Due to the low cell numbers obtained after CD45 depletion only ¼ of the recommended quantity of exploratory antibodies was used. Plates were incubated at 4°C for 30min in the dark. Thereafter, fixation was performed using the Cytofix buffer (BD Biosciences) for 1 hour at 4°C in the dark. As an additional backbone marker, cells were stained intracellularly for anti-AIRE AF750 in Cytoperm buffer (BD Biosciences) and one well stained with anti-FOXN1 (Rode et al., 2015) PE as an additional exploratory marker, over-night at 4°C in the dark. The next day cells were resuspended in 100 μl FACS buffer before analysis.

### Infinity Flow and single-cell clustering and expression analysis

For the Infinity Flow computational analysis of the LEGENDScreen datasets, the acquired fcs files were gated on CD45 negative cells or specifically on EpCAM1^+^ TEC using the FlowJo software. The newly exported fcs files were then used as the dataset for the Infinity Flow pipeline as recently published (Becht et al., 2021). The augmented data matrices generated during this process were then further analysed using the Seurat package for hierarchical clustering of the cells and differential expression analysis (Hao et al., 2021). Genes were filtered by hand to exclude T-cell related and focus on stromal cell related genes (Table S1). Values below zero were set to zero to allow for log normalization.

We compared the Infinity Flow data matrices with the scRNAseq dataset of (Baran-Gale et al., 2020) by identifying the most closely related genes for each Infinity flow protein, e.g. UEA1 fluorescence was identified with Fut1 RNA expression, since FUT1 synthesizes the glycan target of UEA1. Clusters from each dataset were then compared using the SingleR package in R (Aran et al., 2019).

### Histological analyses

Frozen thymus tissue sections (7μm) were fixed in acetone and stained using antibodies specific for CD69 (1:100, H1.2F3, BioLegend), Ly51 (1:200, 6C3, BioLegend), K8 (1:500, TROMA-1, NICHD supported Hybridoma Bank), K14 (1:500, Poly19053, BioLegend). Images were acquired using a Leica DMi8 microscope.

### Reaggregate thymic organ cultures

Perinatal cTEC (CD45^-^EpCAM1^+^MHCII^+^Ly51^+^CD83^+^CD40^+^) and non-perinatal cTEC (CD45^-^EpCAM1^+^MHCII^+^Ly51^+^CD83^-^CD40^-^) were sorted from the thymi of 2-week-old C57BL/6 mice and put in co-cultures with CD69^-^ DP thymocytes sorted from the same thymi, respectively. For this cells were transferred in a 1:1 TEC to DP ratio into 1.5 mL tubes containing 1 mL Iscove’s modified Dulbecco’s medium (IMDM) supplemented with 10% FCS, 100 units/mL penicillin, 100 μg/mL streptomycin and 1x GlutaMAX supplement (Gibco). Co-cultures were maintained at 37°C in a humidified atmosphere containing 10% CO_2_ for 48 hours and then analysed by FACS. As a control DP cells were also cultured without the addition of TEC.

### Anti-CD3 injections

6- to 7-week-old Rag2^-/-^ animals were injected intraperitoneally with 50ug of anti-CD3 (clone KT3) or HBSS. Four weeks post injection thymi were analysed for the appearance of DP thymocytes and for changes within their cTEC compartment.

### Cellular indexing of transcriptomes and epitopes by sequencing (CITEseq)

Cells were isolated from six thymi of 1-week- and three thymi of 16-week-old C57BL/6 mice and depleted of CD45^+^ cells by AutoMACS. Subsequently cells were stained for CD45, EpCAM1, Ly51, Ter119 and with PI. In addition cells were stained with antibodies coupled to oligonucleotides directed against CD9, CD40, CD49a, CD54, CD63, CD73, CD83, CD117, CD146 (human with cross reactivity to mouse), CD200, CD270 (HVEM), CD274, Ly6D, Ly6C/Ly6G (Gr1), MadCAM1, Podoplanin, CD80, CD86, MHCII, Sca1, CD31, EpCAM1, CD36, CD133, CD157, CD300LG (Biolegend, see Table S1), and the *Ulex europaeus* agglutinin I (UEA1) lectin labeled with biotin, followed by secondary staining with streptavidin-PE coupled to an oligonucleotide. CD45^-^Ter119^-^ EpCAM1^+^ and CD45^-^Ter119^-^EpCAM1^-^ cells were sorted in a 70% to 30% ratio into a 1.5 mL tube containing FACS buffer for the 1-week-old and 16-week-old samples, respectively. For both timepoints an estimate of 28000 total cells were loaded on two wells of a 10x Genomics Chromium Single Cell Controller. After single-cell capture cDNA and library preparation were performed according to the manufacturer’s instructions using a Single-Cell 3’ v3 Reagent Kit (10x Genomics) with the changes as described in (Stoeckius et al., 2017) to capture cDNA and produce libraries from antibody derived oligos (ADT). Sequencing was performed on one lane of the Illumina NovaSeq 6000 system with a mix of 90% cDNA library and 10% ADT library resulting in 151nt-long paired-end reads.

The dataset was analysed by the Bioinformatics Core Facility, Department of Biomedicine, University of Basel. cDNA reads were aligned to ‘mm10’ genome using Ensembl 102 gene models with the STARsolo tool (v2.7.9a) with default parameter values except the following parameters: soloUMIlen=12, soloBarcodeReadLength=0, clipAdapterType=CellRanger4, outFilterType=BySJout, outFilterMultimapNmax=10, outSAMmultNmax=1, soloType=CB_UMI_Simple, outFilterScoreMin=30, soloCBmatchWLtype=1MM_multi_Nbase_pseudocounts, soloUMIfiltering=MultiGeneUMI_CR, soloUMIdedup=1MM_CR, soloCellFilter=None. ADT libraries were also processed using the STARsolo tool with default parameters except soloCBmatchWLtype=1MM_multi_Nbase_pseudocounts, soloUMIfiltering= MultiGeneUMI_CR, soloUMIdedup=1MM_CR, soloCellFilter=None, clipAdapterType=False, soloType=CB_UMI_Simple, soloBarcodeReadLength=0, soloUMIlen=12, clip3pNbases=136. Further analysis steps were performed using R (v4.1.2). Note that cell filtering was done based only on the analysis of the gene expression, not ADT abundance. Cells were considered as high-quality cells if they had at least 2000 UMI counts, which is the threshold derived from the distribution of UMI counts across cells, forming a data set of 9953 cells.

Multiple Bioconductor (v3.14) packages including DropletUtils (v1.14.2), scDblFinder (v1.8.0), scran (v1.22.1), scater (v1.22.0), scuttle (1.4.0) and batchelor (v1.10.0) were applied for the further analysis of the data set mostly following the steps of the workflow presented at https://bioconductor.org/books/release/OSCA/. Normalised (Lun et al., 2016) log-count values for the gene expression were used to construct a shared nearest-neighbour graph (Xu and Su, 2015), which nodes, i.e. cells, were clustered by ‘cluster_louvain’ method from the R igraph package (Blondel et al., 2008). Counts reflecting the ADT abundance in cells were also log-normalised and clustered in a similar manner.

The data set was subjected to the cell-type annotation using the Bioconductor package SingleR (v1.8.1) and samples from the Immunological Genome Project (ImmGen) provided by the Bioconducter package celldex (v1.4.0) as the reference. Clusters of cells mostly assigned to ‘Epithelial cells’ (5834 cells) were filtered (Figure S6D). Note that one of the clusters (cluster A, Figure S6A) was excluded at this step, because it was mostly composed of cells with elevated percentage of reads mapping to mitochondrial and ribosomal genes and lower number of counts.

The gene expression of filtered cells was re-analysed by removing the batch effect formed by the combination of the sample of origin and the number of counts per cell (cells with >12000 counts and cells with 12000 counts) and re-clustered (Figure 5A,B). Cells were also subjected to the cell-type annotation using scRNAseq transcriptional profiles of single TEC as the reference data set (Baran-Gale et al., 2020) (Figure 5D,F). The scoreMarkers function of the scran package was applied to find marker genes of clusters 1-3. The standardised log-fold change across all pairwise comparisons ‘mean.logFC.cohen’>1 was used as the significance threshold defining the set of marker genes.

A t-SNE dimensionality reduction was used for visualizing single cells on two dimensions. T-SNE coordinates were calculated using the runTSNE function from the scater package and default parameters. For the visualization of cells based on the gene expression, coordinates of principal components and 2000 most variable genes with excluded mitochondrial and ribosomal genes were used as the input. For the visualization of cells based on the ADT abundance, coordinates of principal components and all ADTs were used as the input.

## Author contributions

F.K., C.V-V., S.M., S.Z., L.M., I.C-A., M.E.D, A.W., and B.L. performed experiments; F.K., C.V-V., S.P., A.B., S.Z., L.M., and J.R. analysed data; F.D. provided *Csnb*^Cre^ mice; G.A. provided the thymi from *Pou2f3*^*-/-*^ mice; F.K. and G.A.H. conceived the project and wrote the manuscript.

## Acknowledgements

We thank Emilie Cosway and Sonia Parnell for their technical assistance and Lilly von Muenchow for critical reading of the manuscript. Further we would like to acknowledge Grozdan Cvijetic for his help to set up the Infinity Flow computational analysis pipeline. We also thank Jakub Abramson for sharing the *Csnb*^Cre^ mice.

This work was supported by the Swiss National Science Foundation (IZLJZ3_171050; 310030_184672), the Wellcome Trust (105045/Z/14/Z) and the National Institute for Health Research (NIHR) Oxford Biomedical Research Centre (BRC) to G.A.H.; Swiss National Science Foundation Early Postdoc.Mobility Fellowship (P2BSP3_188183) and Postdoc.Mobility Fellowship (P500PB_206823) to F.K.; NIHR Clinical Lectureship to F.D.; work in the G.A. lab is supported by an MRC Programme Grant (MR/T029765/1).

