## Supplementary material for "Combined multidimensional single-cell protein and RNA profiling dissects the cellular and functional heterogeneity of thymic epithelial cells": Suppl. Figures 1-8

Figure S1

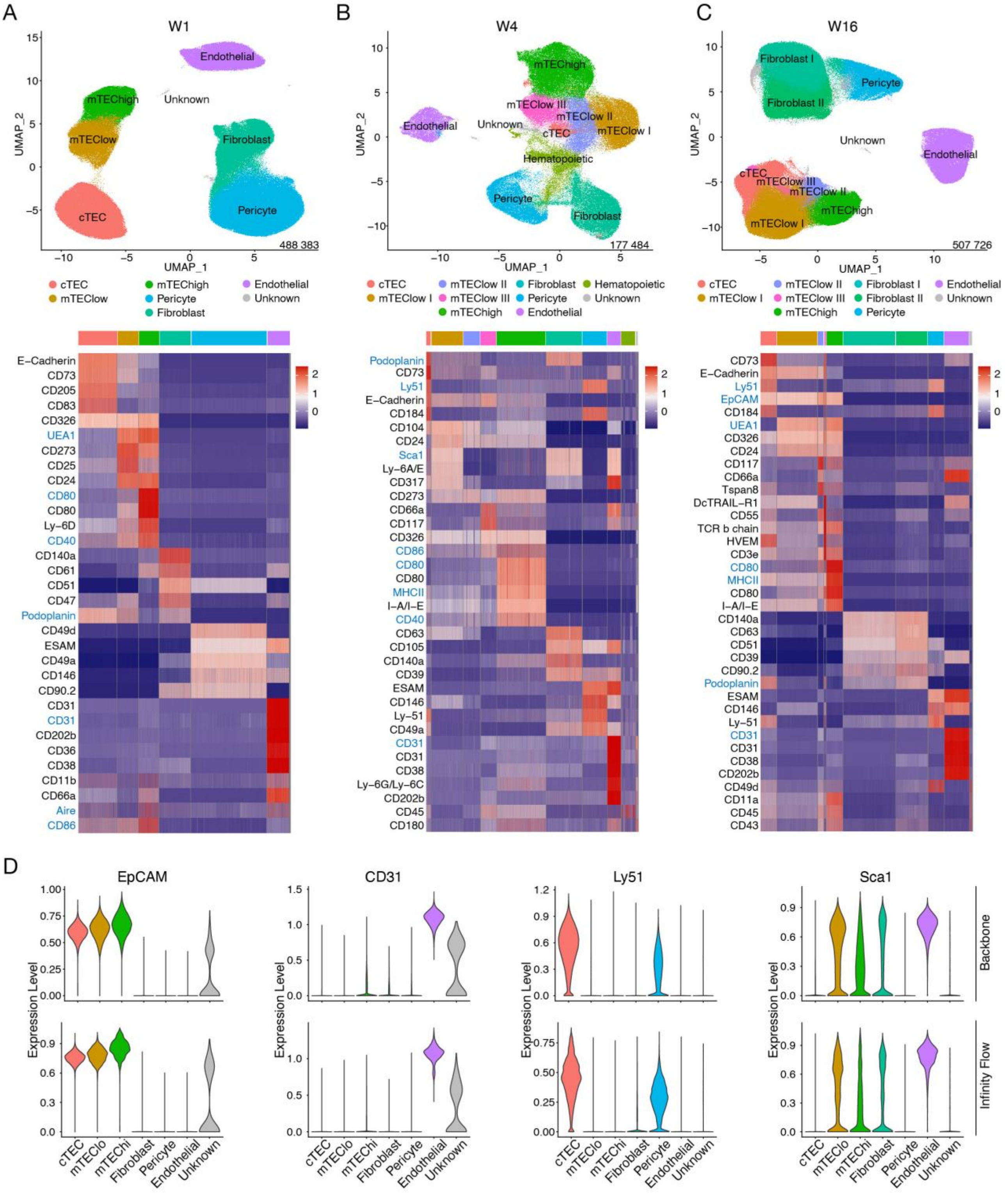


### Figure S1. Infinity Flow analysis on thymic stromal cells

**(A-C)** Infinity Flow analysis was used to impute the expression of surface markers on CD45$^-$ cells derived from thymi of (A) 1-, (B) 4-, and (C) 16-week-old mice. Hierarchical clustering analysis was performed on (A) 488383, (B) 177484, and (C) 507726 CD45$^-$ cells, respectively, and projected in a 2-dimensional space using UMAP (top panels; 7 to 10 clusters were obtained per timepoint). Each colour represents a specific cluster as indicated. Heatmaps (bottom panels) display the expression of the top 7 markers upregulated in each cluster (log fold-change > 0.2). Backbone markers have a blue font. **(D)** Violin plots comparing the expression of the indicated markers based on the backbone staining (top panels) and the prediction of Infinity Flow based on exploratory measurements of the same proteins (bottom panels).

Figure S2

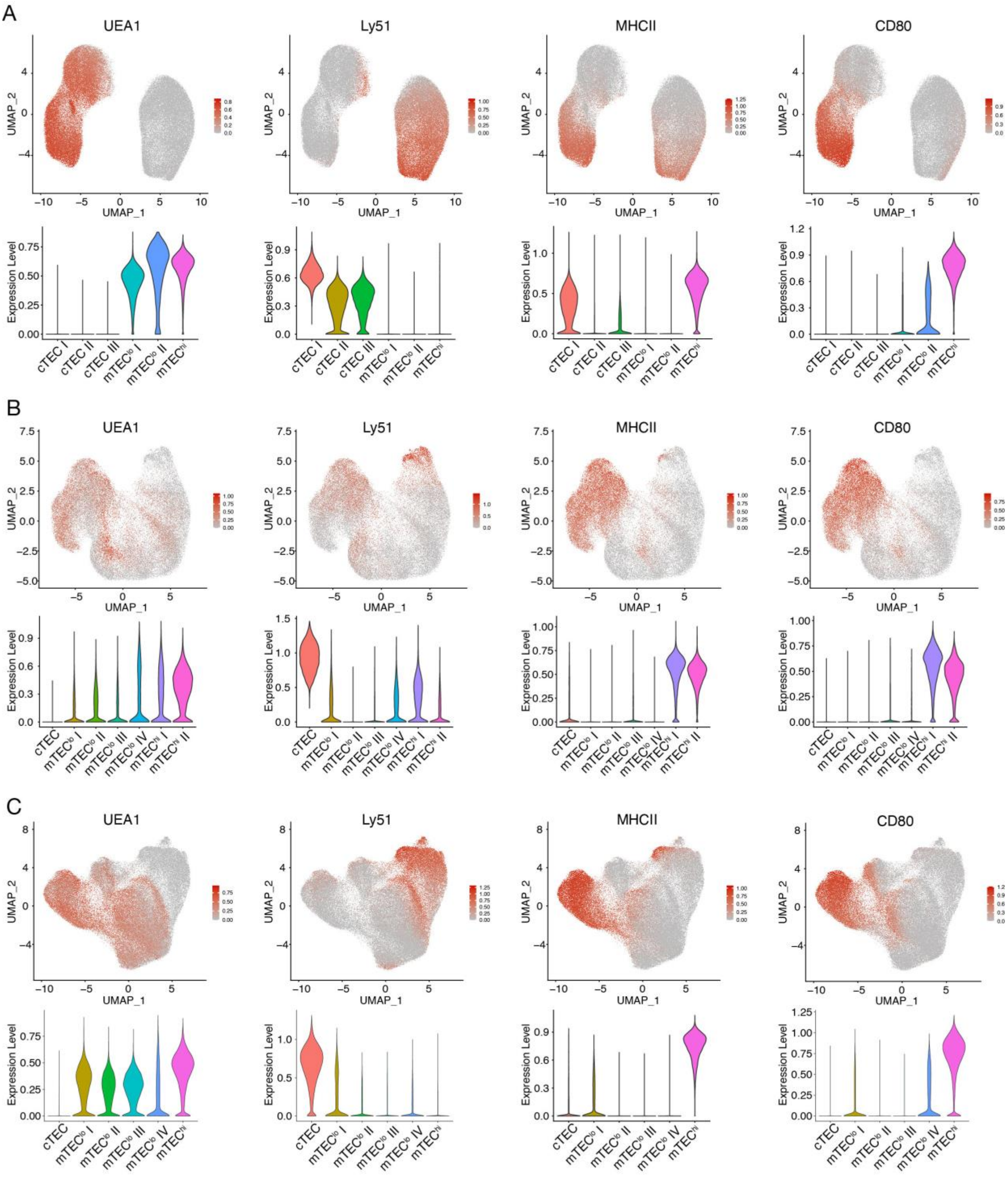


**Figure S2. Classification of clusters based on UEA1, Ly51, MHCII, and CD80 expression**

**(A-C)** UMAP graphs (top panels) and violin plots (bottom panels) illustrating the expression of UEA1, Ly51, MHCII, and CD80 on TEC from (A) 1-, (B) 4-, and (C) 16-week-old mice. Colour gradient indicates expression levels in the UMAP graphs and colours in the violin plots represent the different clusters, as defined in Figure 1B.

Figure S3

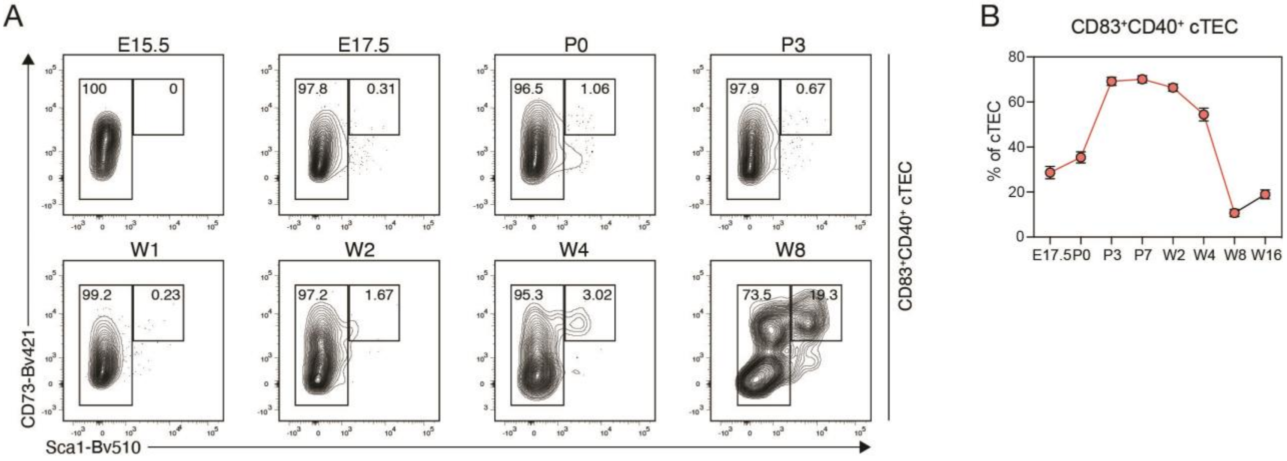


**Figure S3. Characterization of perinatal cTEC**

**(A**) Appearance of a CD73 and Sca1 double positive population within perinatal cTEC was analysed at the indicated timepoints. Shown are representative FACS plots. **(B)** Abundance of a CD83 and CD40 double positive population (hereafter perinatal cTEC) within cTEC was analysed at the indicated timepoints. Shown are cumulative data depicting the percent of perinatal cTEC within cTEC. Data are derived from 2-3 independent experiments per timepoint. Error bars indicate SEM.

Figure S4

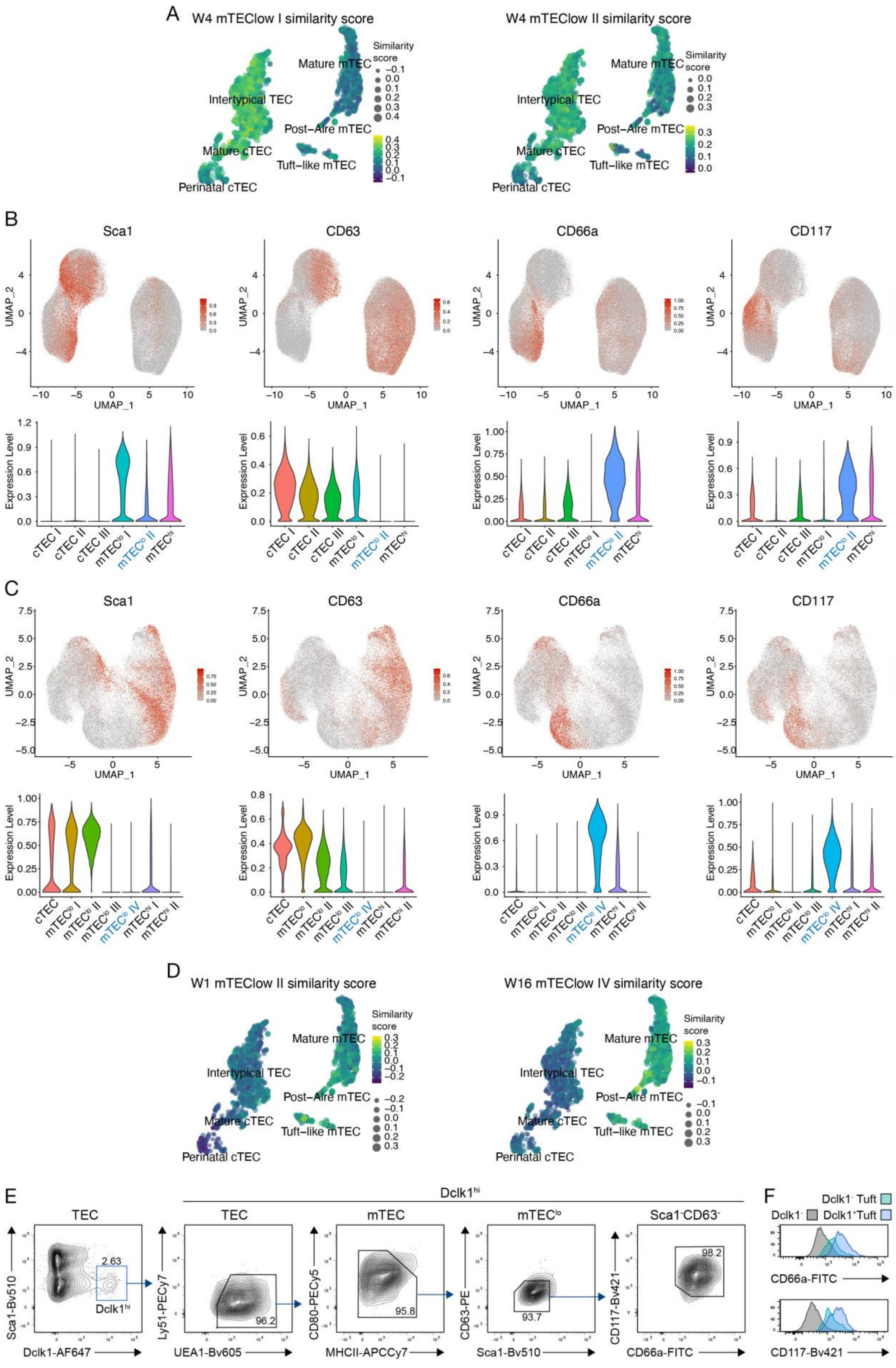
A
W4 mTEClow I similarity score
W4 mTEClow II similarity score
Similarity score
Mature mTEC
Intertypical TEC
Post−Aire mTEC
Mature cTEC
Tuft−like mTEC
Perinatal cTEC
B
Sca1
CD63
CD66a
CD117
UMAP_1
UMAP_2
Expression Level
cTEC I
cTEC II
cTEC III
mTEClo I
mTEClo II
mTEChi
C
cTEC
mTEClo III
mTEClo IV
mTEChi I
mTEChi II
D
W1 mTEClow II similarity score
W16 mTEClow IV similarity score
E
TEC
Dclk1hi
mTEC
mTEClo
Sca1-CD63-
Sca1-Bv510
Dclk1-AF647
Ly51-PECy7
UEA1-Bv605
CD80-PECy5
MHCII-APCCy7
CD63-PE
CD117-Bv421
CD66a-FITC
2.63
96.2
95.8
93.7
98.2
F
Dclk1- Tuft
Dclk1-
Dclk1+Tuft

**Figure S4. Dissecting mTEC$^{lo}$ heterogeneity**

**(A)** UMAP graph illustrating the similarity score of the mTEC$^{lo}$ I and II clusters from the 4-week Infinity Flow datasets to each cell of the single-cell RNA-seq reference dataset, based on the surface protein expression levels imputed by Infinity Flow. **(B,C)** UMAP graphs (top panels) and violin plots (bottom panels) illustrating the expression of Sca1, CD63, CD66a, and CD117 on TEC from (B) 1-, (C) 4-week-old mice. Colour gradient indicates expression levels in the UMAP graphs and colours in the violin plots represent the different clusters, as defined in Figure 1B. **(D)** UMAP graph illustrating the similarity score of the mTEC$^{lo}$ II cluster from the 1- (left panel) and the mTEC$^{lo}$ IV cluster from the 4-week (right panel) Infinity Flow datasets to each cell of the single-cell RNA-seq reference dataset, based on the surface protein expression levels imputed by Infinity Flow. **(E)** FACS plots illustrating the percent of Dclk1$^{+}$ TEC falling within the new tuft-like mTEC gating strategy, as defined in Figure 5D. **(F)** Histograms illustrating the expression levels of CD66a and CD117 within Dclk1$^{-}$ TEC, Dclk1$^{-}$ tuft-like mTEC and Dclk1$^{+}$ tuft-like mTEC.

Figure S5

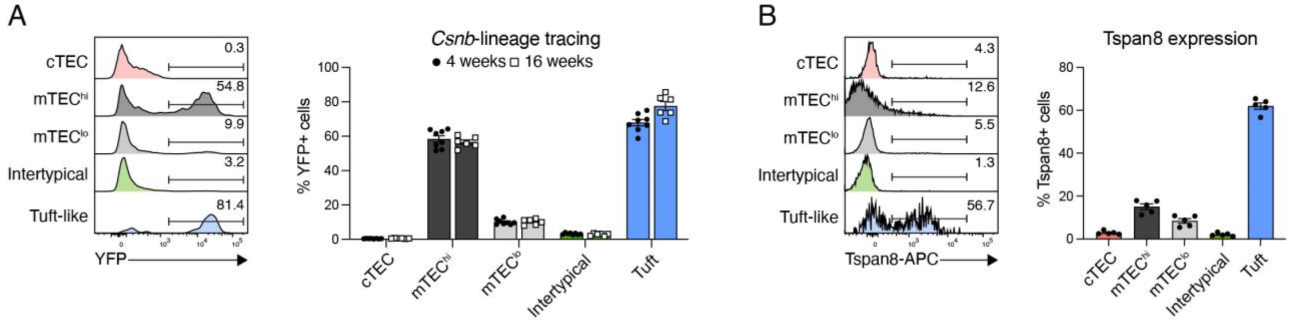


**Figure S5. Pre-mature and post-Aire mTEC compartments**

**(A)** *Csnb*$^{Cre}$::Rosa26$^{LSL\text{-}YFP}$ mice were analysed for the abundance of YFP$^{+}$ cells within cTEC, mTEC$^{hi}$, mTEC$^{lo}$, intertypical TEC and tuft-like mTEC at 4 and 16 weeks after birth. Shown are representative histograms and cumulative data. Data are derived from two independent experiments per timepoint. Error bars indicate SEM. **(B)** WT mice were analysed for the abundance of Tspan8$^{+}$ cells within cTEC, mTEC$^{hi}$, mTEC$^{lo}$, intertypical TEC and tuft-like mTEC. Shown are representative histograms and cumulative data. Data are derived from two independent experiments. Error bars indicate SEM.

Figure S6

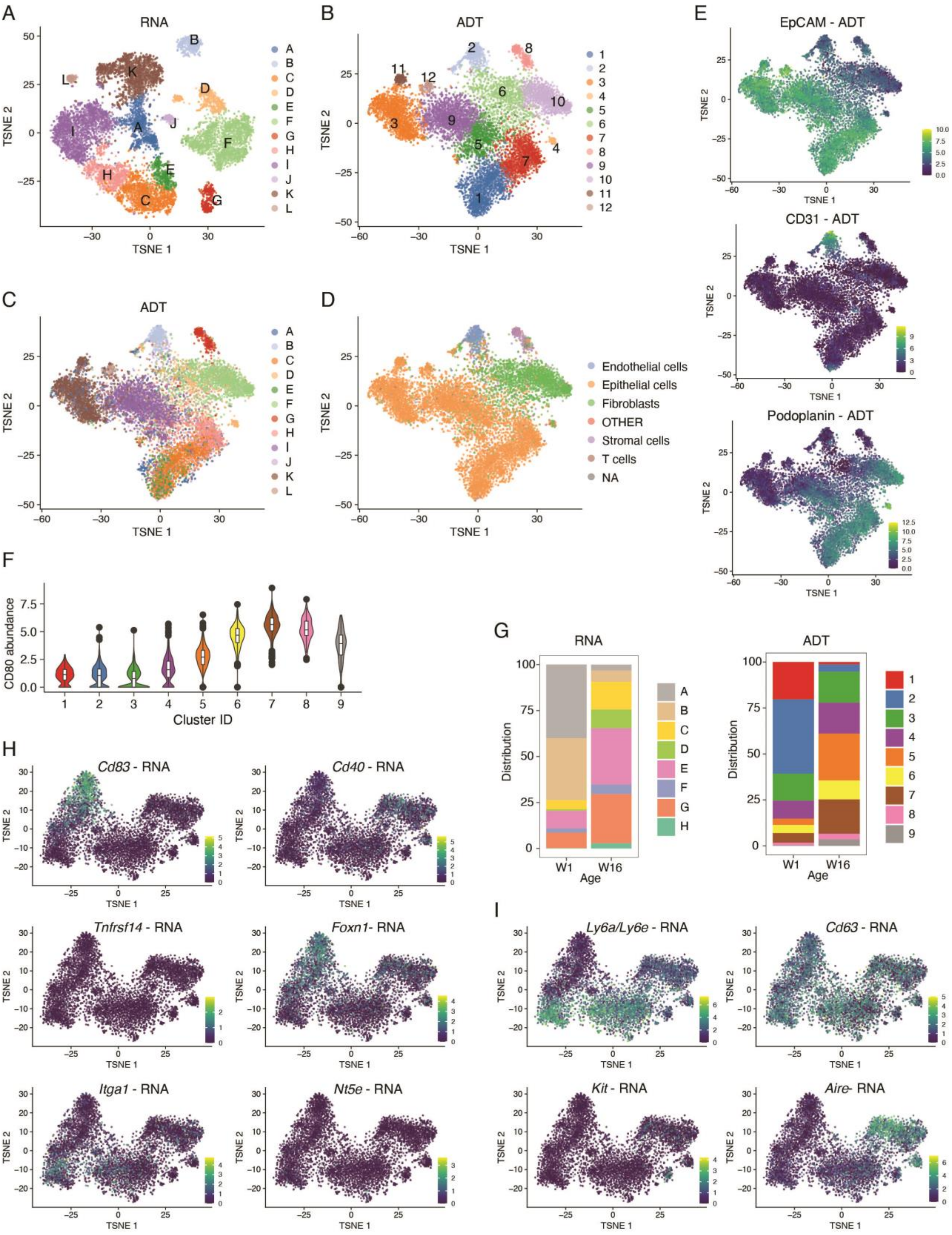


**Figure S6. CITEseq analysis on thymic stromal cells**

CD45$^-$Ter119$^-$ thymic stromal cells isolated from 1- and 16-week-old WT mice were used for scRNAseq in combination with CITEseq as described in the methods. **(A-C)** Hierarchical clustering analysis was performed on 9953 cells either using (A) the gene expression analysis or (B) only considering the detection of ADTs. Results were projected in a 2D space using t-SNE. Each colour represents a specific cluster. In (C) t-SNE distribution of the ADT clustering is shown using the cluster colouring of the RNA analysis. **(D)** Cells were annotated based on transcriptional similarity to reference datasets derived from the Immunological Genome Project (ImmGen). Each colour represents a specific subset as defined in the reference dataset. **(E)** T-SNE plots illustrating the scaled expression of EpCAM1, CD31, and Podoplanin across ADT clusters. **(F)** Violin plots depicting the abundance of CD80 ADTs across ADT TEC clusters. **(G)** Bar graphs illustrating the distribution of 1- and 16-week-old derived TEC across ADT (left panel) and RNA (right panel) clusters. Each colour represents a specific cluster. **(H,I)** T-SNE plots illustrating the scaled expression of the gene expression of (H) perinatal cTEC markers such as *Cd83*, *Cd40*, *Tnfrsf14*, *Foxn1*, *Itga1*, and *Nt5e*, and of (H) tuft-like and intertypical TEC markers such as *Ly6a/Ly6e*, *Cd63*, *Kit*, and *Aire* across ADT clusters.

Figure S7

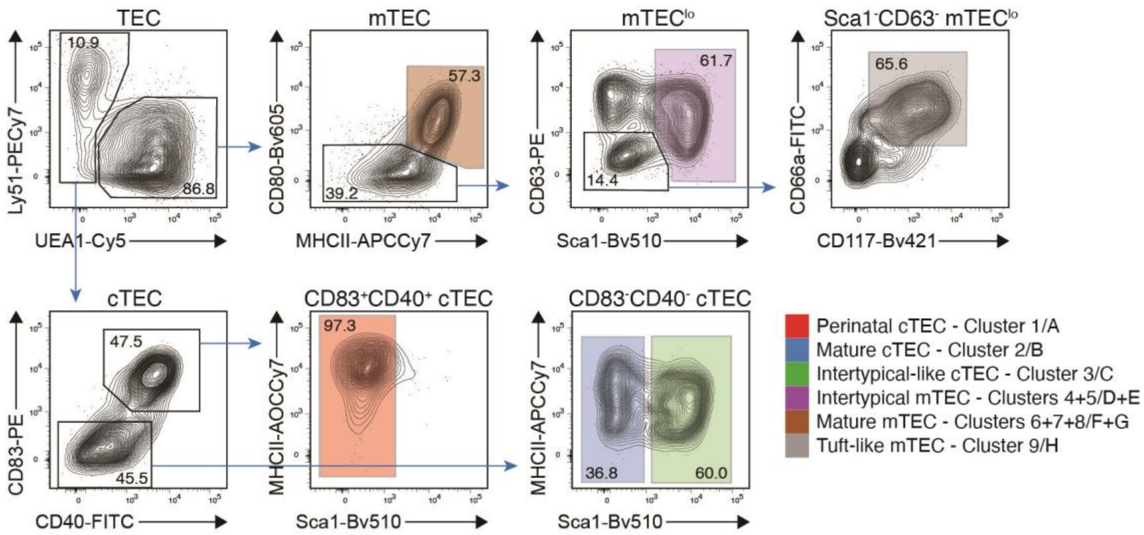


**Figure S7. Gating strategy to identify TEC subpopulations**

Shown are representative FACS plots illustrating the new gating strategy to identify TEC subpopulations based on the surface expression profiles obtained from CITEseq. Colours represent different TEC subpopulations and CITEseq clusters as indicated. Data are derived from a 4-week-old WT mouse and is representative of 3 independent experiments.

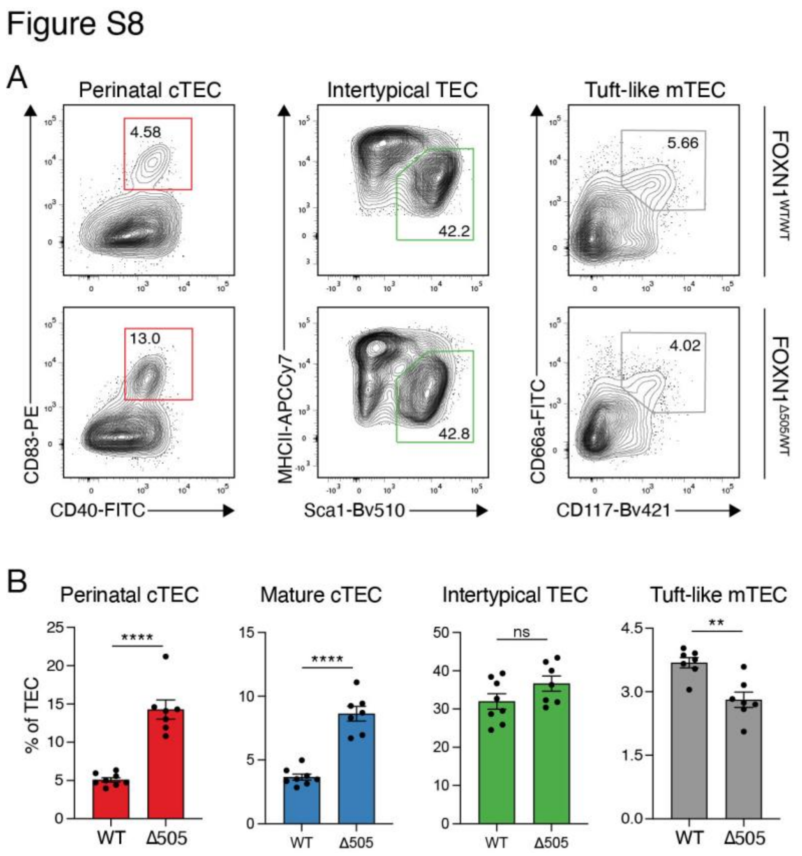


**Figure S8. Application of new TEC markers on FOXN1$^{\Delta505/WT}$ mice**

**(A,B)** FOXN1$^{\Delta505/WT}$ mice were analysed for the abundance of perinatal cTEC, mature cTEC, intertypical TEC and tuft-like mTEC using the new markers and compared to WT mice at an age of 4-weeks. Shown are (A) representative FACS plots and (B) cumulative data. Data are derived from three independent experiments. Error bars indicate SEM. Statistical analysis was done with two-tailed unpaired Student`s t-test. **, $P < 0.01$; ****, $P < 0.0001$.
